# Improved two-stage GWAS identifies rare genetic variants associated with dermo susceptibility in the eastern oyster

**DOI:** 10.64898/2026.09.08.750289

**Authors:** Paul Coyne, Henry Sun, Zhenwei Wang, Sandra Casas, Jerome La Peyre, Mason Williams, Scott Rikard, David Bushek, Ximing Guo

## Abstract

*Perkinsus marinus* is a protist that causes the wasting disease dermo in the Eastern oyster, *Crassostrea virginica*. The disease causes high mortality of both wild and cultured oysters, impacting ecosystem health and aquaculture. Despite decades of research, the genetic mechanisms of dermo resistance remain poorly understood. Traditional selective breeding based on phenotypes has yielded improvements in dermo resistance, but progress has been slow, likely due to the trait’s complex genetic architecture. Understanding the genetic mechanisms underlying dermo resistance may facilitate advanced breeding to accelerate improvement. This study conducted genome-wide association studies (GWAS) of dermo resistance, leveraging a multi-generational dataset comprising 2,423 oysters phenotyped for survival following a dermo challenge and genotyped using a high-density 66K SNP array. A two-stage GWAS with more inclusive quality control identified 48 dermo-resistance markers, including many associated with genes for innate immune response and energy metabolism. Most dermo-resistance markers showed rare minor alleles occurring at higher frequencies in oysters that died after challenge, suggesting that these rare alleles are deleterious and associated with dermo susceptibility. The strongest association was with a polymorphism in the *mucin-5AC-like* gene, explaining 8.1% of phenotypic variation and suggesting that mucus production plays a crucial role in dermo resistance. The inclusion of these markers has improved genomic prediction accuracy, and their identification provides new insights into genomic variation and the architecture of dermo resistance, informing future strategies for genetic improvement.

**ARTICLE SUMMARY:** Dermo disease, caused by the protist *Perkinsus marinus*, is a major threat to Eastern oyster (*Crassostrea virginica*) aquaculture and reef restoration. This study analyzed dermo resistance in 2,423 oysters genotyped with a 66k SNP array. Forty-eight putative dermo-resistance markers were identified, including many within genes involved in mucus production, energy metabolism, and immune response. Most dermo-resistance markers were loci carrying rare alleles associated with susceptibility, suggesting that rare genetic variants play a critical role in shaping fitness-related traits. These findings help reveal genetic mechanisms underlying dermo resistance and provide new breeding strategies to support oyster aquaculture.

## INTRODUCTION

Eastern oysters (*Crassostrea virginica*) support a major U.S. aquaculture industry, generating more than $195 million in commercial landings in 2024 (NOAA Fisheries 2024). Throughout their native range, from the Gulf of St. Lawrence to the Gulf of Mexico, oysters are exposed to *Perkinsus marinus*, the protist pathogen that causes dermo disease (Bower 2025, RSSBP 2026). In some locations, yearly dermo-mortality rate can exceed 50% for market-sized (>75 mm) oysters (Burreson and Ragone Calvo 1996, Soniat 1996); the establishment of dermo disease in Delaware Bay doubled the annual background mortality rate on moderately infested oyster beds and quadrupled it on heavily infested beds (Bushek et al. 2012). Transmission of the parasite is direct as infectious propagules released into surrounding waters from live and dead oysters are filtered by living oysters (Bushek et al. 2002, Ragone Calvo et al. 2003, Audemard et al. 2006, Allam et al. 2013, Lau et al. 2018, Gignoux-Wolfsohn et al. 2021). The parasites initially infect epithelial tissues and are readily ingested by hemocytes which spread infections to other tissues (Mackin and Boswell 1956, La Peyre et al 1995, Tasumi and Vasta 2007, Lau et al 2018, Carnegie et al 2021). As the number of parasites increases, inflammation, characterized by an increase in hemocyte density, is observed, followed by tissue damage and eventually death of the emaciated oysters (Mackin 1951, Ray et al 1953).

The rate at which dermo progresses is influenced by both environmental and genetic factors. The parasite proliferates most actively in warmer (>20°C) and more saline (>12 ppt) waters, resulting in summer and early fall oyster mortalities (Burreson & Ragone Calvo 1996, Soniat 1996). It has been demonstrated that there are several strains of *P. marinus* with varying levels of virulence (Bushek and Allen 1996), and there are also *C. virginica* populations in dermo-endemic regions exhibiting differing levels of resistance to the pathogen (Bushek and Allen 1996, Brown et al. 2005, Casas et al. 2017, La Peyre et al. 2019), suggesting a heritable component to dermo resistance.

Traditional selective breeding for dermo resistance is a key priority of many Eastern oyster breeding programs, but progress has been slow. The selected stocks remain partly susceptible to the disease and are often followed by unexplained decreases in survival and higher dermo infection intensities in later generations (Ragone Calvo et al. 2003, Degremont et al. 2015, Casas et al. 2017, Guo 2021). This is likely due to several factors such as low heritability of resistance, a complex genetic architecture, and a lack of consistent selection pressure, all of which lead to low selection efficiency (Wang et al. 2025). Further, traditional selective breeding relies on survival to a marketable size as the phenotype, which takes 2-3 years to measure in field-deployed oysters. Advanced approaches such as genomic selection and targeted modifications may accelerate the development of dermo-resistant stocks.

The sequencing of the Eastern oyster genome and the development of tools such as high-density single-nucleotide polymorphism (SNP) arrays have enabled advanced genomic analyses of important traits and genomic approaches to genetic improvement (Gómez-Chiarri et al. 2015, Guo et al. 2023, Puritz et al. 2024). Wang et al. (2025) demonstrated that genomic selection is more effective than phenotypic selection in breeding for dermo resistance. While genomic selection uses genome-wide SNPs to predict breeding values and does not require prior knowledge of marker-trait association, the inclusion of known disease-resistance markers may improve the accuracy of genomic prediction and selection. Despite many studies of genes and pathways that respond to *P. marinus* infection, few genetic markers have been identified that confer dermo resistance (Yu and Guo 2006, Tasumi and Vasta 2007, Wang et al. 2010, He et al. 2012, Soudant et al. 2013, Proestou and Sullivan 2020, Chan et al. 2021). Disease-resistance markers are best identified through genome-wide association studies (GWAS) (Uffelmann et al. 2021). Using GWAS, Wang et al (2025) identified 32 loci with modest associations to dermo resistance, although none were significant after adjustment for multiple testing, likely due to the limited sample size (n = 508) or statistical power. Another study, however, using an order-of-magnitude larger sample size (n = 5,720), recently identified six markers significantly associated with dermo resistance (Delomas et al. 2026).

In this study, a GWAS of dermo resistance was conducted using 2,423 oysters genotyped with the OysterCv 66K SNP array (Guo et al. 2023). The genotyped oysters came from training populations for genomic selection over three generations, where the oysters either died or survived lab-based challenges with *P. marinus*. By combining multiple generations of data to increase statistical power and using a two-stage GWAS with inclusive data quality control (QC), this study identified 48 SNPs significantly associated with dermo resistance, along with associated genes involved in host defense. The inclusion of these markers has significantly improved the accuracy of genomic prediction of dermo resistance (Sun et al. 2026), indicating that these markers and associated genes provide strong candidates for targeted selection or for the development of dedicated marker panels to improve dermo resistance.

## MATERIALS AND METHODS

### Oysters and Challenge Experiments

The Eastern oysters used in this study were from three successive generations (F0-F2) of training populations used for genomic selection for dermo resistance (Morris et al. 2026). Oysters of the training population were challenged with *P. marinus* using established protocols (Wang et al. 2025). The F0 generation was collected from a wild population in St. Andrew Bay, Florida, under the Florida Fish and Wildlife Conservation Commission special activity license number SAL-22–2391-SR. The F0 oysters were challenged in 2022 by injection of *P. marinus* into the shell cavity, with details of the full challenge available in Wang et al. 2025. Oysters in the challenge experiment were monitored until cumulative mortality reached approximately 50%. Tissue from live (resistant) and dead (susceptible) oysters was collected and genotyped with the OysterCv 66K SNP array (Guo et al. 2023). Genomic selection models were trained from an F0 training population and used to predict genomic estimated breeding values (GEBVs) in a breeding population of unchallenged oysters from the same wild population. The F0 oysters with the highest GEBVs were selected to produce the F1 genomic selected line (FLGS).

Survivors from the F0 challenge were selected to produce the F1 phenotypic selected line (FLP). In 2023, equal numbers of oysters from F1 FLGS and FLP lines were selected and subjected to the same challenge (except for parasite inocula; 1 M parasites/g tissue was used to challenge F0 oysters versus 5 M parasites/g tissue for F1 and F2 oysters) to create the F1 training population, and the same genomic selection protocol was later used to produce the F2 generation. In 2024, the same process was repeated with the F2 generation to produce the F3 generation. The three training populations (live and dead) created by challenging F0, F1 and F2 oysters were combined and used for this study. The combined dataset consisted of 2,423 oysters across the three generations, including 243 live and 262 dead from F0, 519 live and 421 dead from F1 and 494 live and 484 dead from F2.

### SNP Genotyping and Data Quality Control

Mantle tissue was dissected and fixed in 100% ethanol from oysters immediately following mortality or, for survivors, once overall mortality reached about 50%. Survival following challenge was recorded as a binary trait (live or dead). Time-to-death was also recorded and analyzed; however, because it yielded results consistent with but not as strong as the binary survival analysis, only the latter results are presented.

All oysters were genotyped with the 66K OysterCv SNP array using the Affymetrix GeneTitan instrument through a service provider (Guo et al. 2023). SNPs were called using the Axiom Analysis Suite software (ThermoFisher, CA), following the Best and Recommended Workflow. Individuals with genotype call rates (CRs), defined as the proportion of assayed SNPs for which a genotype was successfully assigned, below 90% were excluded. SNP call rate was defined as the proportion of individuals for which a genotype was successfully assigned at a given SNP. To evaluate the influence of marker filtering on GWAS performance, several QC strategies were examined, varying in minor allele frequency (MAF) and SNP CR thresholds. Standard QC filtering required MAF > 0.05 and SNP CR > 95%, whereas expanded analyses relaxed the MAF threshold to MAF > 0.01 or 0.005. These alternative QC strategies allowed assessment of the tradeoff between retaining informative rare variants and minimizing false-positive associations.

A two-stage GWAS strategy was used to identify and evaluate dermo-resistance markers (DRMs). First, an exploratory GWAS was performed to identify candidate dermo-resistance loci, with all SNP markers retained after removing individuals with genotype CRs <90%, without additional filtering based on MAF or SNP CR. Markers exceeding the chromosome-wide significance threshold (defined below) and having a SNP CR > 0.85 were designated candidate DRMs. Candidate DRMs with SNP CRs between 0.85 and 0.95 and/or MAFs between 0.01 and 0.05 were considered conditional DRMs and evaluated further based on allele- and genotype-frequency patterns across the three generations. Markers showing consistent patterns expected for true resistance-associated loci were retained for subsequent analyses despite failing conventional filtering criteria. Prior to all GWAS, any remaining missing genotypes were imputed using a k-nearest neighbors strategy (k = 10) applied separately within each generation (Troyanskaya et al. 2001).

### Genome-wide Association Analysis

Genome-wide association analyses were performed using the ASRgwas package in R in tandem with the ASReml-R package to estimate genomic variance components and fit GWAS models (Butler et al. 2023). Before association testing, genomic relationship matrices and principal components were generated using the pre.gwas() function to account for population structure and relatedness among individuals. Association testing per SNP was conducted using linear mixed models implemented in gwas.asreml() using the formula [*y* = *Xβ* + *ma* + *Zu* + *ɛ*], where X is the fixed-effects matrix corrected for population structure, *β* are the estimated coefficients for X, *m* is the genotype of a given individual, ɑ is the additive genetic effects of a given SNP, *Z* is an individual identity matrix, *u* accounts for the polygenic effects of SNPs not being tested, and *ε* is residual error. Based on an inspection of the principal component distributions, seven principal components were included as fixed effects, along with the intercept. Both Gaussian and binomial models yielded comparable association results; therefore, the Gaussian model was used throughout due to its lower computational requirements. The significance of marker-trait associations was evaluated using both genome-wide and chromosome-wide Bonferroni-adjusted p-value thresholds. The genome-wide threshold was calculated as *p* < (0.05 / number of analyzed SNPs), and the chromosome-wide threshold was calculated as *p* < (0.05 / number of analyzed SNPs) × number of chromosomes, yielding results comparable to linkage-disequilibrium-adjusted significance thresholds while remaining straightforward to reproduce.

Five additional GWAS were conducted using markers from different QC strategies following the initial exploratory GWAS used to identify candidate markers (Table 1). These marker QC strategies progressively lowered the MAF threshold and incorporated conditional DRMs that would have been removed by standard QC to assess whether including rare variants and conditional DRMs improved model fit and the detection of significant associations. GWAS performance and fit were assessed using the genomic inflation factor (λ) and quantile-quantile (QQ) plots. The genomic inflation factor was calculated as the ratio of the observed median chi-square test statistic to its expected median under the null hypothesis of no association (Devlin and Roeder 1999). Consequently, λ provides a measure of whether the distribution of test statistics is inflated because of population stratification, cryptic relatedness, or other systematic biases. Values of λ approaching 1.0 were considered indicative of appropriate control of population stratification, whereas substantial deviations suggested either inflation or overcorrection of association statistics. Because λ is calculated from the median GWAS test statistic, its sensitivity to a relatively small number of highly significant loci is limited. Therefore, λ was used primarily to evaluate overall model calibration and the effects of the broader QC strategy rather than to justify retention of individual DRMs. QQ plots were inspected to assess the deviation of the observed test-statistic distribution from the expected null distribution.

**Table 1.** Different QC strategies for minor allele frequency (MAF) and call rate (CR) used for extended GWAS to identify dermo-resistance markers (DRMs). Asterisks denote cases where conditional DRMs with lower CRs (0.85-0.95) were retained despite failing traditional QC.

| QC Strategy | MAF | SNP CR | Sample CR | Purpose |
| --- | --- | --- | --- | --- |
| Exploratory | None | None | > 0.90 | Exploratory GWAS used to identify candidate DRMs. |
| Traditional | >0.05 | >0.95 | >0.90 | Standard GWAS quality-control parameters |
| Inclusive | > 0.01 | > 0.95 | > 0.90 | Relaxed MAF threshold to include SNPs with rare alleles |
| Traditional Plus | >0.05 | >0.95* | >0.90 | Standard QC plus conditional DRMs |
| Inclusive Plus | > 0.01 | > 0.95* | > 0.90 | Relaxed MAF threshold plus conditional DRMs |
| Most Inclusive Plus | > 0.005 | > 0.95* | > 0.90 | Most permissive MAF threshold plus conditional DRMs |

Narrow-sense heritability was estimated from the variance components of fitted linear mixed models using residual maximum likelihood (REML) (Corbeil and Searle 1976). Heritability (*h*^2^) was defined as 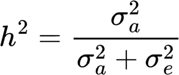 where 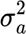 is the additive genetic variance captured by all autosomal SNPs and 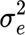 is the residual variance (Rameez et al. 2022). QQ and Manhattan plots to visualize GWAS results were generated using qqman (Turner 2014).

### Functional Analysis of Dermo-Resistance Markers

Markers significantly associated with dermo resistance at the chromosome-level threshold were mapped to the *C. virginica* reference genome (GCA_002022765.2) using positions defined by the 66k OysterCv SNP array. All variants were classified as exonic, intronic, or intergenic. For intergenic variants, the nearest annotated gene was assigned based on physical proximity, as linkage disequilibrium (LD) may extend to genes around the SNP (Sved and Hill 2018). Protein annotations for gene products were obtained from UniProt (UniProt Consortium, 2025). For genes lacking curated protein annotations, predicted protein sequences were queried against the non-redundant protein database using BLASTX (e-value < 1 × 10^-10^) (Altschul et al., 1990). Gene Ontology (GO) terms for annotated genes were retrieved from UniProt when available. A total of 61.4% of genes across the GCA_002022765.2 assembly, 48.0% of genes represented on the SNP array, and 29.1% of DRMs lacked Biological Process GO term annotations. For proteins lacking GO annotations, BLASTP searches against the Swiss- Prot database (Boeckmann et al. 2003) were performed using homologous proteins from well-annotated species, primarily *Homo sapiens*, to infer putative biological functions. Because sequence homology does not necessarily imply conservation of function across distantly related taxa, functions inferred from homologs were considered putative and interpreted cautiously.

Functional assignments for implicated genes were manually curated following recommendations for annotation of non-model organisms (Primmer et al. 2013). GO terms for identified DRMs were grouped into broad functional categories adapted from the generic GO Slim framework and manually modified to better represent molluscan physiology, with classification groups presented in Supplemental Table 1 (Gene Ontology Consortium 2000). Due to the limited number of background genes with GO terms for *C. virginica*, a generic additive ranking calculation was used to summarize the abundance and significance of each cluster of GO terms rather than conventional enrichment tests (modified from the Pathway scoring algorithm in Lamparter et al. 2016). The cluster scoring metric is defined as 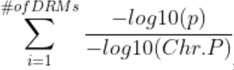, where *P* is the p-value for a given SNP and *Chr.P* is the chromosome-wide p-value threshold. Each DRM contributed a maximum of one score to each functional category, regardless of the number of GO terms assigned to that marker.

## RESULTS

### Genome-wide Association Analysis

After removing individuals with CR < 0.90 (n = 66), 2,357 individuals remained for downstream analyses, including 493 F0, 900 F1, and 964 F2 oysters, of which 53.3% survived the dermo challenge. The number of SNPs retained after different QC strategies is given in Table 2. On average, SNP CRs are slightly lower in oysters that died than in survivors (96.8% versus 98.2%), possibly because of DNA degradation in deceased individuals.

**Table 2.** Performance metrics of GWAS with marker sets after different strategies of marker QC, including SNPs retained, genomic inflation factor (λ), genome-wide p-value threshold (Gen. P) and genome-wide DRM (Gen. DRM) identified, chromosome-wide p- value threshold (Chr. P) and chromosome-wide DRM (Chr. DRM) identified, conditional DRMs at relaxed threshold recovered (cDRM Rec’d), false-discovery rate (FDR), heritability (h2) and its standard error (h2 SE).

| QC | SNPs | $\lambda$ | Gen. P | Gen. DRM | Chr. P | Chr. DRM | cDRM Rec'd | FDR | $h^2$ | $h^2$ SE |
| --- | --- | --- | --- | --- | --- | --- | --- | --- | --- | --- |
| Exploratory | 65,860 | 1.031 | 7.59E-07 | 32 | 7.59E-06 | 44 | 31 | 0.160 | 0.316 | 0.038 |
| Traditional | 39,923 | 0.994 | 1.25E-06 | 0 | 1.25E-05 | 2 | 0 | N/A | 0.238 | 0.037 |
| Inclusive | 48,139 | 1.004 | 1.04E-06 | 15 | 1.04E-05 | 24 | 10 | 0.342 | 0.253 | 0.037 |
| Traditional Plus | 39,940 | 0.989 | 1.25E-06 | 15 | 1.25E-05 | 17 | 16 | 0.342 | 0.248 | 0.037 |
| Inclusive Plus | 48,160 | 1.009 | 1.04E-06 | 35 | 1.04E-05 | 48 | 31 | 0.143 | 0.262 | 0.037 |
| Most Inclusive Plus | 50,301 | 1.013 | 9.94E-07 | 41 | 9.94E-06 | 50 | 31 | 0.125 | 0.264 | 0.037 |

The exploratory GWAS with all SNPs identified 44 markers exceeding the chromosome-wide significance threshold, of which 31 were conditional DRMs with substandard CRs (0.85 - 0.95) and MAFs (0.01 - 0.05) (Table 2). Among these conditional DRMs, 21 had SNP CRs below the traditional 0.95 threshold and 15 had MAF <0.05, indicating that all 31 loci would have been removed from the dataset under traditional or standard QC. None of the DRMs had MAF values below 0.01. Sixteen of the 31 DRMs also exceeded the genome-wide significance threshold. As expected, the exploratory GWAS exhibited an inflated genomic inflation factor (λ), suggesting that the later implemented marker QC and optimization were necessary.

After more stringent marker QC, GWAS substantially improved model fit, as supported by λ values approaching 1.0 and corresponding QQ plots. GWAS with markers after traditional QC of MAF > 0.05 and CR > 0.95, had a genomic inflation factor λ = 0.994, indicating minimal inflation, but identified only two significant DRMs at the chromosome-wide threshold. This result indicates that the traditional QC threshold is too stringent to detect all DRMs. On the other hand, GWAS with the marker set from the most inclusive QC threshold (MAF > 0.005, CR > 0.95, plus the 31 conditional DRMs identified at relaxed QC) produced the largest number of DRMs (50) at the chromosome-wide significance level (Table 2). However, it also produced a higher λ, suggesting inflated test statistics and an increased likelihood of false positives.

Among the evaluated QC strategies, markers from the Inclusive Plus protocol (MAF > 0.01, CR > 0.95, plus the 31 conditional DRMs identified at the relaxed threshold) provided the best overall performance, combining acceptable genomic inflation (λ = 1.009), the second-lowest false discovery rate, and the second-greatest number of significant DRMs (Table 2). In addition to recovering the 31 conditional DRMs, this marker set identified 17 additional DRMs, 15 more than the marker set from traditional QC, bringing the total number of DRMs to 48. The QQ and Manhattan plots from GWAS with this marker set are presented in Figure 1. While no strong peaks were observed on the Manhattan plot, there was some clustering of DRMs on regions of chromosomes 1, 3, and 9.

**Figure 1.**
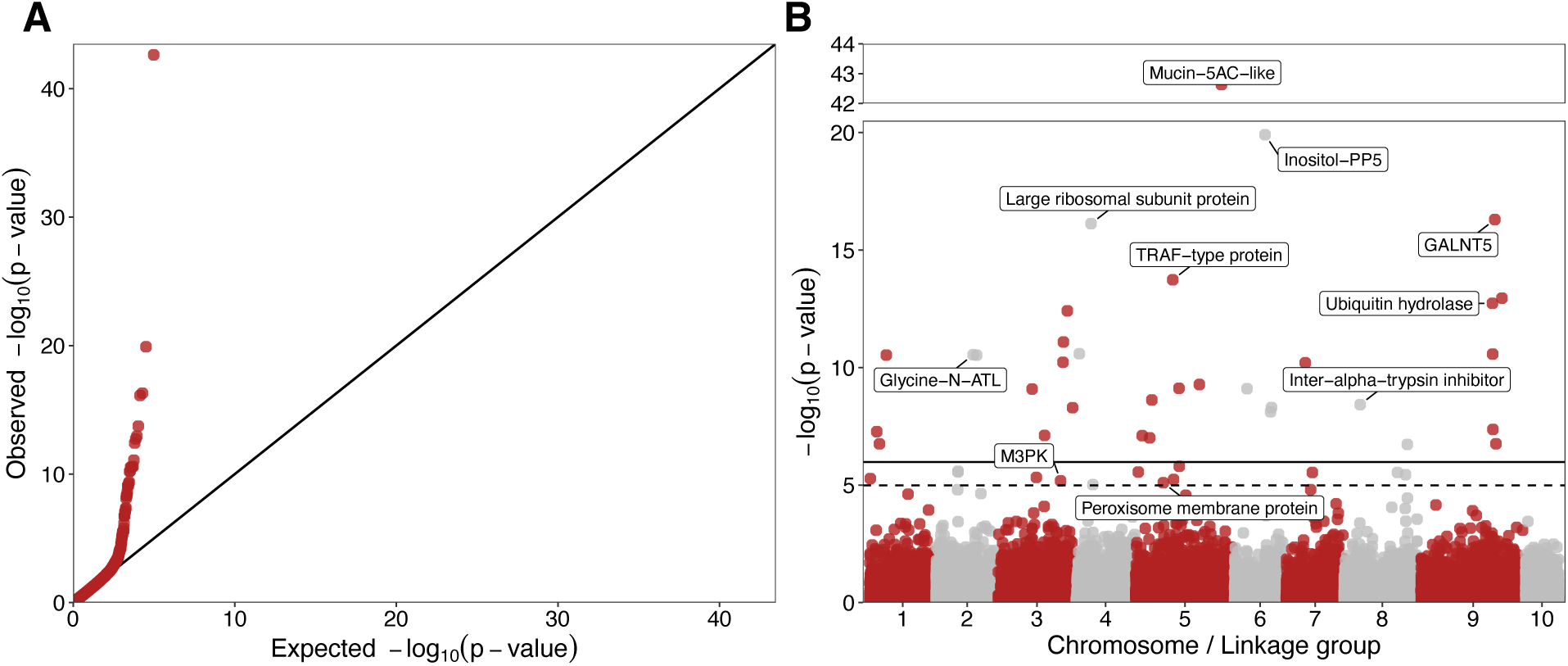
GWAS results using 48,160 SNPs from the Inclusive Plus QC strategy. (A) QQ plot assessing GWAS model fit. (B) Bifurcated Manhattan plot showing loci with significant associations with dermo resistance at genome (solid line) and chromosome (dash line) level thresholds. Several genes of high significance and/or relevant functional annotations are noted.

The estimated narrow-sense heritability varied among different marker sets. The unfiltered marker set yielded the highest estimate, likely inflated. Estimates using marker sets from different QC strategies were mostly consistent, ranging from 0.238 to 0.264 (Table 2). The selected Inclusive Plus model estimated the heritability of dermo resistance to be 0.262 ± 0.037. The inclusion of the 31 conditional DRMs with lower CRs (0.85 - 0.95) increased heritability estimates by ∼0.01.

### Characterization of Dermo-Resistance Markers

The DRMs identified in this study tended to exhibit low polymorphism. Of the 48 DRMs, 32 (67%) had MAF ≤ 0.05, and 41 (85%) DRMs had a MAF ≤ 0.10 (Table 3). Only 7 of the 48 DRMs had MAF > 0.10, and none had MAF > 0.18. Of the 48,160 SNPs used, SNPs with MAF ≤ 0.05 accounted for only 17%, SNPs with MAF ≤ 0.10 accounted for 29%, and SNPs with MAF ≥ 0.18 accounted for 49%. This result suggests that DRMs tend to be loci with rare alleles. At all but one DRMs, the MAF is higher in dead oysters than in oysters that survived the dermo challenge (Figure 2; Supplementary Data File S1), suggesting that the minor allele is more susceptible to dermo than the major allele. It should be noted that the SNP CRs in this study are lower in dead than in live oysters, likely due to poor DNA quality from tissue degradation. DNA quality is known to affect call rate, but it cannot account for the higher MAF observed in dead oysters. Poor DNA quality may cause allelic dropout in heterozygotes, thereby underestimating MAF.

**Figure 2.**
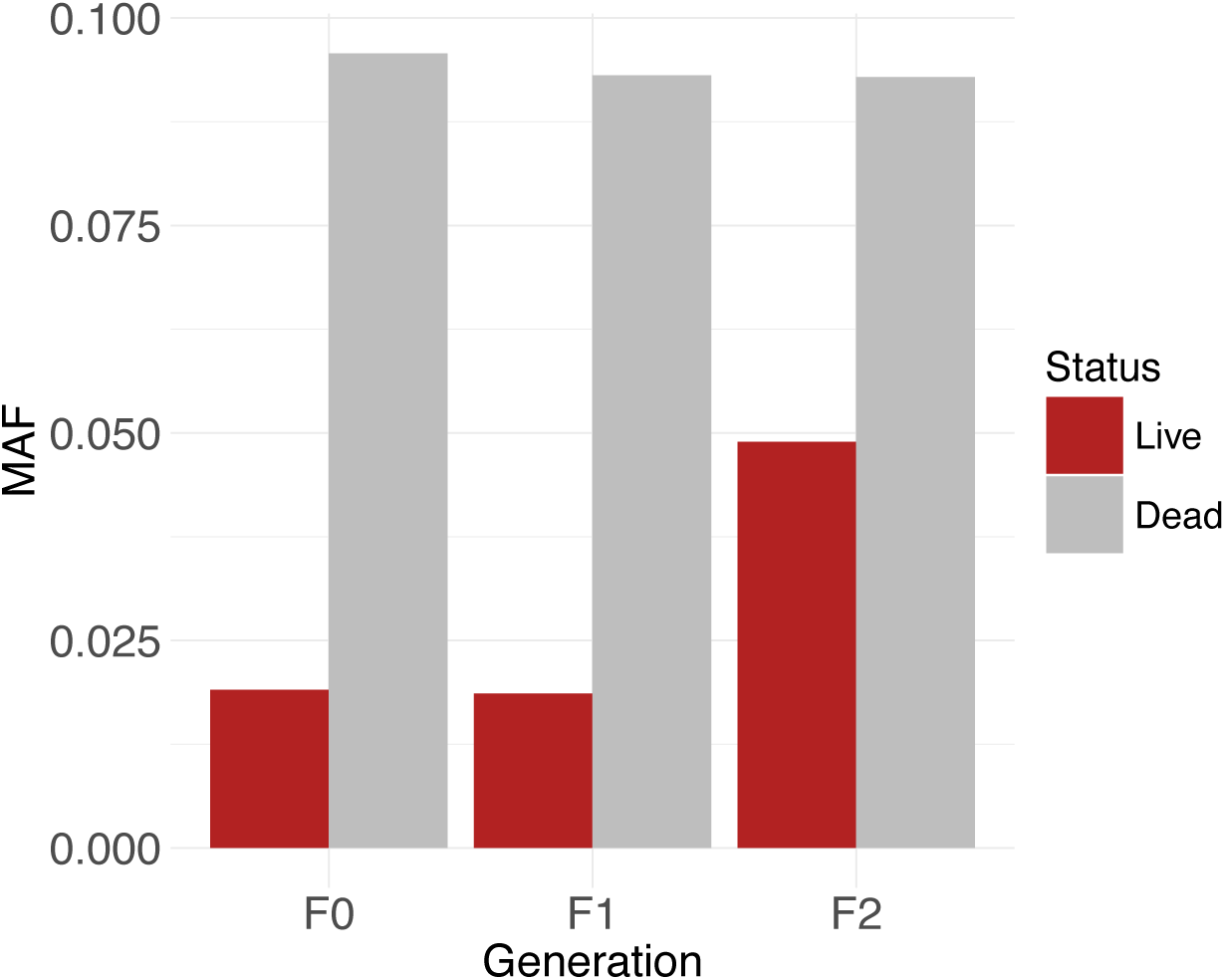
Minor-allele frequency in live and dead oysters at the 48 DRMs identified by GWAS, showing consistently higher frequency in dead oysters across three generations.

**Table 3.** Dermo-resistance markers and associated genes identified by GWAS. For each SNP marker, the chromosome (Chr) position, call rate (CR), minor allele frequency (MAF), p-value and variance explained (VarEx), SNP type (S-type), geneID and description of the associated gene are given. SNPs marked with * in the SNP marker column denote non-synonymous mutations. SNPs found in the non-coding (NC) region have their proximity to the nearest coding region noted in kilobase pairs (kb).

| SNP marker | Chr | Position | CR | MAF | p-Value | VarEx | S-type | GeneID | Gene Description |
| --- | --- | --- | --- | --- | --- | --- | --- | --- | --- |
| AX-563280343 | 5 | 85,778,717 | 0.85 | 0.10 | 2.40E-43 | 8.1 | Exon | LOC111133191 | Mucin-5AC-like |
| AX-564181349 | 6 | 30,498,160 | 0.92 | 0.03 | 1.20E-20 | 3.5 | Intron | LOC111100314 | inositol-polyphosphate 5-phosphatase |
| AX-564439151 | 9 | 73,582,137 | 0.93 | 0.05 | 5.00E-17 | 2.5 | UTR | LOC111112794 | Polypeptide N-acetylgalactosaminyltransferase |
| AX-563911687 | 4 | 15,162,551 | 0.87 | 0.15 | 7.50E-17 | 4.9 | Exon | LOC111131333 | Large ribosomal subunit protein, 39S, L49 |
| AX-570458254 | 5 | 37,453,443 | 0.98 | 0.02 | 1.80E-14 | 2.3 | Exon | LOC111136201 | TRAF-type zinc finger domain-containing protein 1-like |
| AX-576810971 | 9 | 80,692,085 | 0.99 | 0.02 | 1.10E-13 | 2.2 | Intron | LOC111114934 | Uncharacterized protein |
| AX-567945231 | 9 | 71,177,090 | 0.88 | 0.07 | 1.90E-13 | 1.2 | Intron | LOC111112808 | Ubiquitin carboxyl-terminal hydrolase 40-like |
| AX-563866890 | 3 | 68,720,351 | 0.98 | 0.02 | 3.80E-13 | 2.0 | Exon | LOC111124407 | Uncharacterized protein |
| AX-563859187 | 3 | 64,829,703 | 0.92 | 0.10 | 8.20E-12 | 4.3 | Exon | LOC111126109 | Nucleoside diphosphate kinase |
| AX-570229026 | 4 | 3,487,124 | 0.93 | 0.04 | 2.60E-11 | 1.6 | Intron | LOC111127877 | Centrosomal protein of 170 kDa-like |
| AX-574123461 | 9 | 71,483,352 | 0.92 | 0.10 | 2.70E-11 | 3.5 | NC, 1kb | LOC111116130 | Coiled-coil domain-containing protein |
| AX-563453997 | 1 | 16,453,540 | 0.95 | 0.03 | 2.90E-11 | 1.5 | NC, 58kb | LOC111122823 | Diacylglycerol kinase |
| AX-574557543 | 2 | 37,071,622 | 0.99 | 0.01 | 2.90E-11 | 1.6 | NC, 2kb | LOC111118592 | Glycine N-acyltransferase-like protein |
| AX-563703370 | 2 | 40,490,953 | 0.99 | 0.02 | 3.00E-11 | 1.7 | NC, <1kb | LOC111119180 | Uncharacterized protein |
| AX-562975825 | 3 | 64,336,735 | 0.92 | 0.11 | 6.00E-11 | 3.5 | Exon | LOC111125816 | Uncharacterized protein |
| AX-570890079 | 7 | 19,262,204 | 0.86 | 0.14 | 6.30E-11 | 2.6 | Intron | LOC111102434 | Protein lin-28 homolog |
| AX-576402328 | 7 | 36,966,661 | 0.91 | 0.02 | 4.20E-10 | 1.5 | Intron | LOC111104376 | Monocarboxylate transporter 12-like |
| AX-564118820 | 5 | 63,888,812 | 0.98 | 0.01 | 5.30E-10 | 1.4 | Intron | LOC111134722 | Collagen alpha-1(IV) chain-like |
| AX-564085983 | 5 | 43,629,915 | 0.90 | 0.13 | 7.60E-10 | 3.8 | NC, <1kb | LOC111099077 | Uncharacterized protein |
| AX-563298068 | 6 | 12,610,178 | 0.92 | 0.17 | 7.90E-10 | 3.7 | UTR | LOC111099734 | Uncharacterized protein |
| AX-574116952 | 3 | 33,752,651 | 0.99 | 0.04 | 8.20E-10 | 1.2 | NC, <1kb | LOC111122494 | Uncharacterized protein |
| AX-568235396* | 5 | 16,766,320 | 0.93 | 0.04 | 2.40E-09 | 1.3 | Exon | LOC111134771 | CCCTC-binding factor |
| AX-576902130 | 8 | 15,904,596 | 0.88 | 0.05 | 3.80E-09 | 1.4 | Intron | LOC111108691 | Inter-alpha-trypsin inhibitor heavy chain H3-like |
| AX-563318564 | 6 | 36,867,607 | 0.89 | 0.10 | 5.00E-09 | 3.6 | Intron | LOC111100897 | Cytosolic purine 5'-nucleotidase-like |
| AX-567276575* | 3 | 73,834,663 | 0.99 | 0.01 | 5.10E-09 | 1.2 | Exon | LOC111123612 | Uncharacterized protein |
| AX-564193914 | 6 | 35,961,286 | 0.99 | 0.02 | 7.70E-09 | 1.0 | Exon | LOC111099479 | MORN repeat-containing protein 4-like |
| AX-574389243 | 1 | 7,032,094 | 0.86 | 0.10 | 5.30E-08 | 2.2 | Intron | LOC111102414 | ATP synthase subunit b |
| AX-570642792 | 5 | 7,160,185 | 0.99 | 0.01 | 6.60E-08 | 1.4 | Exon | LOC111132431 | Uncharacterized protein |
| AX-567116016 | 3 | 46,061,031 | 0.96 | 0.05 | 7.70E-08 | 1.0 | NC, 1kb | LOC111126306 | BarH-like 1 homeobox protein |
| AX-576895722 | 3 | 48,975,738 | 0.97 | 0.06 | 8.40E-08 | 2.0 | UTR | LOC111126881 | Protein Wnt |
| AX-570332532 | 5 | 14,590,782 | 0.87 | 0.14 | 9.80E-08 | 2.3 | Intron | LOC111133995 | Sex peptide receptor-like |
| AX-563564797 | 9 | 71,752,745 | 0.98 | 0.01 | 1.30E-07 | 1.2 | Intron | LOC111114595 | HECT-type E3 ubiquitin transferase |
| AX-574408724 | 1 | 9,706,268 | 0.99 | 0.01 | 3.30E-07 | 1.2 | Intron | LOC111127446 | Zinc finger protein GLI4-like |
| AX-567965988 | 9 | 74,759,147 | 0.99 | 0.01 | 5.60E-07 | 1.2 | Exon | LOC111111318 | D-3-phosphoglycerate dehydrogenase |
| AX-576610402 | 8 | 62,672,745 | 0.99 | 0.01 | 5.90E-07 | 1.2 | NC, 6kb | LOC111109272 | Uncharacterized protein |
| AX-567679698 | 8 | 60,948,169 | 0.99 | 0.01 | 8.50E-07 | 1.0 | NC, 28kb | LOC111108439 | Plexin-B-like |
| AX-570494413* | 5 | 43,967,204 | 0.89 | 0.07 | 1.60E-06 | 1.7 | Exon | LOC111133385 | Protein FAM221A |
| AX-571052524 | 7 | 26,340,103 | 0.98 | 0.01 | 1.70E-06 | 1.1 | UTR | LOC111104804 | Uncharacterized protein |
| AX-574115859 | 2 | 22,018,321 | 0.98 | 0.01 | 2.60E-06 | 1.2 | Exon | LOC111119192 | Angiopoietin-1 receptor-like |
| AX-575695122 | 5 | 3,265,336 | 0.91 | 0.14 | 2.80E-06 | 3.9 | Exon | LOC111136401 | Adenosine receptor A3-like |
| AX-575122210 | 3 | 37,922,336 | 0.99 | 0.04 | 4.50E-06 | 1.1 | Exon | LOC111126324 | Gastrula zinc finger protein XICGF57.1-like |
| AX-574092264 | 2 | 21,959,068 | 0.96 | 0.06 | 5.20E-06 | 0.8 | Intron | LOC111119193 | Pancreatic triacylglycerol lipase-like |
| AX-575653763 | 1 | 598,105 | 0.99 | 0.01 | 5.30E-06 | 0.9 | NC, 3kb | LOC111138315 | R3H domain-containing protein 1-like |
| AX-576896226 | 3 | 61,996,152 | 0.98 | 0.01 | 6.60E-06 | 0.9 | Intron | LOC111124479 | Mitogen-activated protein kinase kinase kinase |
| AX-570465218 | 5 | 38,331,253 | 0.99 | 0.02 | 7.10E-06 | 0.9 | NC, 6kb | LOC111099129 | Uncharacterized protein |
| AX-564057487 | 5 | 28,060,768 | 0.98 | 0.02 | 8.00E-06 | 0.9 | Exon | LOC111135112 | Peroxisomal membrane protein 2-like |
| AX-563408780 | 8 | 52,915,763 | 0.96 | 0.03 | 8.40E-06 | 0.9 | Intron | LOC111106495 | Extensin-like |
| AX-575684868 | 2 | 44,672,366 | 0.98 | 0.02 | 9.10E-06 | 1.1 | Intron | LOC111117952 | Phosphatase and actin regulator 4B-like |

Of the 48 significant SNPs or DRMs identified using SNPs from the Inclusive Plus protocol, 37 were genic and 11 were intergenic (Table 3). Of the genic SNPs, 17 were intronic and 16 were exonic, including 13 synonymous and 3 non-synonymous mutations, and 4 were from the untranslated region. Among the three non-synonymous SNPs, AX-568235396 in the *CCTC-binding factor* gene was a particularly penalizing mutation, as it causes the substitution of valine (an uncharged, nonpolar amino acid) for glutamic acid (a negatively charged, polar amino acid) in a highly conserved zinc-finger motif.

Several DRM-associated genes have well-defined functions in immune and stress response. The strongest association was observed for a synonymous SNP within a *mucin-5AC- like* gene (p = 2.4E-43), which explained approximately 8.1% of the phenotypic variance for dermo resistance (Table 3). The SNP showed consistently higher MAF in dead oysters, indicating that the major allele is associated with dermo resistance (Figure 3A; Supplementary Data File S1). Most of the AA genotypes were found in live oysters, while most of the BB genotypes were found in dead oysters (Figure 3B; Supplementary Data File S1). A SNP in the *mucin-17-like* gene (LOC111107098) also showed consistent MAF difference between live and dead oysters, although the p-value of the association (p = 0.00047) did not reach the chromosome-wide significance threshold. Another highly significant marker was in GALNT5, which encodes a polypeptide N-acetylgalactosaminyltransferase involved in O-linked glycosylation of secreted glycoproteins such as mucin. These results suggest that mucin and related processing genes play an important role in innate immune response and resistance against dermo.

**Figure 3.**
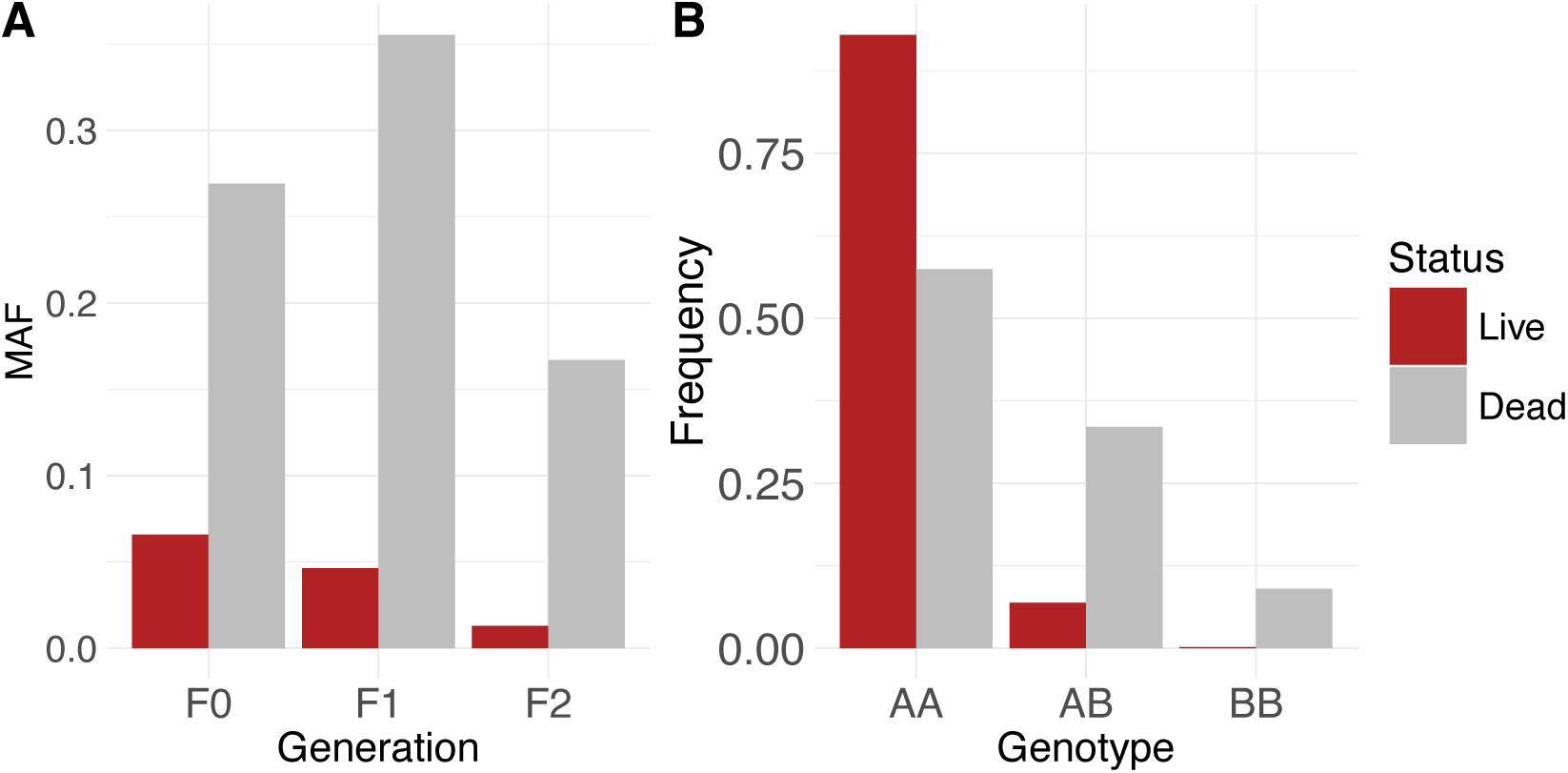
Minor-allele and genotype frequency distribution of SNP AX-563280343 within mucin-5AC-like gene in live and dead Eastern oysters after dermo challenge. (A) Multigenerational trends show consistently higher MAF in dead/susceptible oysters. (B) Genotype frequencies of the SNP in live and dead oysters.

Other implicated genes involved in immune and stress response included 1) *TRAF-type zinc finger domain-containing protein 1-like*, which encodes a signaling adaptor protein that participates in innate immune pathways and stress responses; 2) *Ubiquitin carboxyl-terminal hydrolase 40-like* and *HECT-type E3 ubiquitin transferase*, which may regulate ubiquitination for pathogen degradation; 3) *MORN repeat-containing protein 4-like* and *Plexin-B-like* genes that may regulate phagocytosis; 4) *Protein FAM221A* that may regulate immune response through interaction with the NF-κB pathway; and 5) *Mitogen-activated protein kinase kinase kinase (MAP3K)*, a master regulator of the MAPK signaling for immune and stress response. Also implicated are *collagen alpha-1(IV) chain-like*, *inter-alpha-trypsin inhibitor heavy chain H3-like* and *extensin-like* genes, which may participate in the maintenance and repair of the extracellular matrix.

Several DRMs were found in genes related to metabolism and energy production (Table 3). They included *pancreatic triacylglycerol lipase-like, D-3-phosphoglycerate dehydrogenase, glycine N-acyltransferase-like, cytosolic purine 5’-nucleotidase-like, nucleoside diphosphate kinase, large ribosomal subunit protein 39S L49*, and *ATP synthase subunit b*. Implicated genes also included *protein Wnt* and *BarH-like 1 homeobox* that are known to regulate development and differentiation. Many other implicated genes encode uncharacterized proteins whose functions are currently unknown.

A total of 43/48 (89.5%) of protein-coding genes associated with DRMs had GO terms successfully assigned using a combination of homology-based functional inference and database-derived annotations. The highest functional clustering score for categories of GO terms was in immune functions (19.69), with 12 DRMs mapping to 21 GO terms (Figure 4). Additional highly represented functional categories included cell signaling (11.40; 15 DRMs, 29 GO terms), neurosensory (8.62; 14 DRMs, 33 GO terms), transcription and RNA regulation (7.63; 10 DRMs, 19 GO terms), and metabolism (6.03; 8 DRMs, 31 GO terms). Other lower-ranking functional clusters included biomineralization, cell structure/motility, cell development, mitochondrial and energy, and metabolism (Figure 4). Together, these annotations indicate that DRM-associated genes span a broad range of biological functional categories, indicating that dermo resistance has a complex architecture.

**Figure 4.**
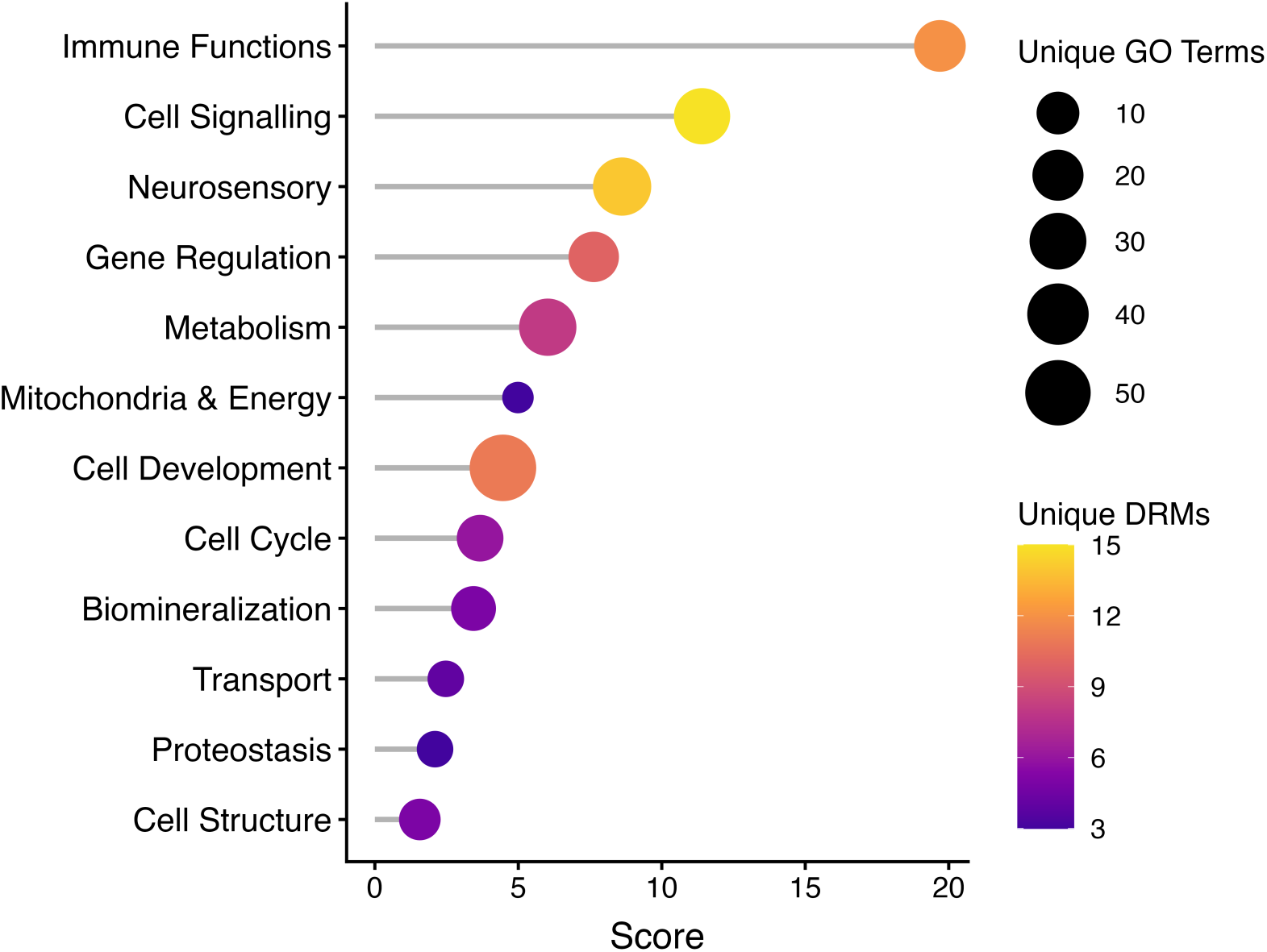
GO terms from each DRM were assigned to broad functional groups, which were then scored based on the p-values of DRMs contributing to each cluster through the equation 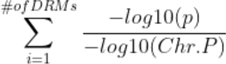, where *P* is the p-value of each individual DRM, *Chr.P* is the chromosome-wide p-value threshold, and each DRM only contributes once per unique cluster. Clustering methodology was manually adjusted from the generic GO slim classification to better reflect bivalve physiology. The size of lollipops signifies the number of unique GO terms identified in each cluster, and the color corresponds to the number of DRMs participating in each cluster.

## DISCUSSION

### New Data QC Approach Improves GWAS

Conventional data QC for GWAS typically removes markers with MAF < 0.05, SNP CRs < 95%, and individual genotype CRs < 90%, with lower MAF thresholds generally reserved for very large populations (>10,000 individuals) (Marees et al. 2018), a sample size not readily available for non-model organisms such as oysters. Applying these conventional QC criteria to the present dataset, either within individual generations or across the pooled population, identified only two chromosome-wide significant loci. This result is consistent with previous studies in the Eastern oyster that identified no or only six significant dermo-resistance markers (Wang et al. 2025, Delomas et al. 2026).

Using a larger number of oysters from three successive generations provided an opportunity to explore new QC approaches for GWAS in this study. The large sample size made it feasible to lower the MAF threshold and include more markers with rare alleles. To improve detection power while limiting false-positive associations, a two-stage GWAS strategy was implemented. An initial exploratory GWAS with all markers identified a set of candidate DRMs, including many that did not meet the conventional QC standards (MAF ≥ 0.05, CR ≥ 0.95), but had MAF ≥ 0.01 and CRs ≥ 0.85. All 31 candidate DRMs identified would be excluded by conventional QC. Closer inspection indicated that nearly all candidate DRMs consistently exhibited differential allele and genotype frequency distributions between live and dead oysters across three generations, as expected for real DRMs. Additionally, most of the DRMs exhibited a dosage effect of alleles on survival. This approach substantially increased the number of biologically supported associations while maintaining acceptable genomic inflation and false discovery rates.

The multigenerational dataset provided an opportunity to evaluate the identified DRMs. Because each generation was partially derived from the previous one, true dermo-resistance loci were expected to exhibit the same differential distribution of alleles and genotypes between surviving and deceased oysters across generations. The observed consistency among nearly all validated DRMs (Supplementary Data File S1) provided greater confidence that these loci represent genuine biological associations rather than statistical artifacts or genotyping errors.

Additional evidence for the biological relevance of DRMs identified herein comes from genomic prediction analyses. All nine genomic prediction models trained on the dataset described in this study showed significant increases in prediction accuracy upon the inclusion of DRMs (Sun et al. 2026). Conversely, other genomic selection studies that relied on conventional SNP filtering techniques observed insignificant improvements in prediction accuracy after increasing SNP marker density (Wang et al. 2025, Delomas et al. 2026), confirming that simply adding more markers does not in itself boost predictive performance for dermo resistance if biologically informative variants remain excluded. The success of this new approach is exemplified in the identification of AX-563280343 in the *mucin-5AC-like* gene, the most significant DRM explaining 8.1% of phenotypic variation. This marker exhibited consistent differences between live and dead oysters across three generations.

### Most Dermo-Resistance Markers are Loci with Rare Alleles

Relaxing the MAF threshold to 0.01 provided an opportunity to interrogate rare genetic variants. The findings that most DRMs were loci with rare alleles, and that the DRMs consistently show higher MAF in dead oysters, suggest that minor rare alleles make oysters more susceptible to dermo. This is supported by the fact that homozygotes for the minor allele (BB) had higher mortality than heterozygotes (AB), who in turn had higher mortality than homozygotes for the major allele (AA). Thus, minor or rare alleles were either directly deleterious or in LD with deleterious mutations, as is well documented in studies of human disease (Dickson et al. 2010, Johansen et al. 2010, Gibson 2012). If this hypothesis is true, the development of dermo resistance would require the gradual purging of these rare deleterious alleles, which would in turn require consistently high dermo pressure in the field.

The persistence of these rare, deleterious alleles may be explained by several evolutionary mechanisms. In dermo-endemic regions, disease pressure varies considerably between years due to environmental and ecological conditions and typically reaches mass-mortality levels only among neighboring reefs (Powell et al. 2012, Kachmar et al. 2025). Spatial variation of dermo pressure across salinity gradients may allow for the preservation of these rare alleles. Populations of oysters in areas of low salinity, with limited resistance to dermo due to reduced selection pressure, may experience gene flow with adjacent, higher-salinity areas experiencing varying dermo pressure (Bushek et al. 2012). Consequently, purifying selection against deleterious alleles is likely intermittent rather than continuous, allowing many potentially deleterious rare variants to persist at low frequencies. Additionally, because dermo resistance appears to be a highly polygenic trait, with susceptibility determined by the cumulative effects of many small-effect loci, the fitness cost of any single rare deleterious allele is likely too small to be efficiently removed by selection (Harrang et al. 2013, Cvijović et al. 2018). Instead, these variants may accumulate at low frequencies while collectively contributing to disease susceptibility (Kryukov et al. 2007).

Additional mechanisms may contribute to the persistence of deleterious rare alleles. First, the unfavorable SNPs may confer or be in linkage with alleles with fitness advantages such as increasing larval survival, resulting in antagonistic pleiotropic effects across the life cycle (Ernande et al. 2003, Yu and Guo 2006, Carter and Nguyen 2011). Secondly, ‘sweepstakes’ reproduction characteristic of broadcast-spawning marine organisms, where a small number of successful parents can contribute disproportionately to each generation, may allow mildly deleterious alleles to persist within the population due to increased genetic drift despite reducing adult fitness (Hedgecock and Pudovkin 2011, Harrang et al. 2013).

The oyster genome is highly polymorphic and contains tens of millions of SNPs, the vast majority of which have a minor allele frequency < 0.05 (Plough et al. 2016, Puritz et al. 2024). The high polymorphism in oysters is likely maintained by balancing selection in adaptation to their biphasic life history and the dynamic environments they face (Guo et al. 2018, Guo 2026a). This study suggests that many rare variants are deleterious under certain conditions, and their existence is part of the population genetic equilibrium maintained by mutation, selection (e.g., balancing selection), genetic drift, and migration (Haldane 1937, Lenz et al. 2016, Yin and Hedgecock 2021, Marion and Noor 2023). This hypothesis is likely applicable to all organisms, especially marine organisms with high fecundity and polymorphism, which have been shown to have a large genetic load with an excess of deleterious mutations (Launey and Hedgecock 2001, Plough 2016). The results of this study highlight the significance of rare genetic variants that have been neglected in most studies. Future studies should attempt to better characterize and identify the primary mechanisms responsible for the maintenance of these susceptibility-associated alleles in natural and selectively bred populations.

### Dermo Resistance is a Polygenic Trait with Low Heritability

Two observations from this study support the notion that dermo resistance is a polygenic trait. First, GWAS revealed no major peaks on the Manhattan plot and no significant loci explaining more than 10% of the variation. The most significant loci explained 8.1% of the variation, and most other DRMs explained 1-4% of the variance. Second, the estimated heritability for dermo resistance, 0.262 ± 0.037 from the best model, is low. This estimate is comparable to the estimate obtained from the F0 generation alone (0.255 ± 0.115) (Wang et al. 2025), and lower than that for several other disease-resistance traits in oysters (Degremont et al. 2015). Selection for resistance to some other oyster diseases, such as MSX (multinucleated sphere X caused by the protozoan *Haplosporidium nelsoni*), ROD (*Roseovarius* oyster disease, caused by the bacterium *Roseovarius crassostreae*) and Ostreid herpesvirus 1 μVar (OsHV-1), was relatively fast, indicating that these diseases may have higher heritability (Guo 2026b). The heritability of 0.262 is lower than the threshold of approximately 0.30 generally considered favorable for efficient phenotypic selection (Goddard et al. 2010), which may be partly responsible for the slow progress in improving dermo resistance through traditional phenotype-based selective breeding. Genomic selection is particularly well suited for polygenic traits with low heritability because it captures the cumulative effects of many loci with small individual effects (Meuwissen et al. 2001). The results of this study therefore support previous findings demonstrating improved prediction accuracy for dermo resistance using genomic selection compared with phenotypic selection (Wang et al. 2025, Delomas et al. 2026).

### Mucus Production is Critical for Dermo Resistance

Functional annotation of significant loci indicates that many underlying determinants of the dermo resistance are components of the innate immune system of Eastern oysters. Approximately 72% of significant markers occurred within or near annotated protein-coding genes, many of which participate in immune function, cellular signaling, transcriptional regulation, apoptosis, or metabolic homeostasis. It should be noted that the function of the many identified genes is inferred from their roles in other organisms but may vary in oysters. The identified SNPs are likely not the causative mutation but are linked to functional variation nearby (Sved and Hill 2018).

The strongest association identified in this study was a synonymous polymorphism within the *mucin-5AC-like* gene (LOC111133191), which explained approximately 8.1% of the observed phenotypic variance. Mucin-5AC is a gel-forming glycoprotein that constitutes a major component of mucus in vertebrates and functions as a primary barrier against invading pathogens (Lang et al. 2007, Wang et al. 2024). Although oyster mucins are structurally less complex than vertebrate homologs, mucus production represents an early line of defense in bivalves by trapping and rejecting pathogens during filter feeding (Allam and Pales Espinosa 2015). This proposed mechanism is supported by previous studies, which demonstrated *P. marinus* mounts a distinct transcriptional response upon exposure to *C. virginica* mucus, indicating that mucus represents a substantial threat to the parasite (Pales Espinosa et al. 2014). A SNP in the *mucin- 17-like* gene is also implicated, although its association (p = 0.00047) did not reach the chromosomal significance threshold (p = 0.0000104). The transcription of *mucin-5AC-like* is upregulated by *P. marinus* infection (Chan et al. 2021). In addition, oyster galectins present within mucus have been shown to specifically recognize *P. marinus*, initiating downstream innate immune responses (Tasumi and Vasta 2007). The identification of genes involved in mucus production in this study strongly supports previous physiological and transcriptomic evidence implicating mucus-mediated pathogen exclusion as a key mechanism of dermo resistance.

Additional support for this hypothesis comes from the association identified near GALNT5, which encodes a polypeptide N-acetylgalactosaminyltransferase responsible for initiating O-linked glycosylation of secreted glycoproteins (Bennett et al. 2012). In vertebrates, GALNT5 contributes to the maturation, stability, and trafficking of mucins, including MUC5AC and MUC1 (Tian et al. 2023, Rapp et al. 2024), suggesting that variation at this locus may indirectly influence dermo resistance by altering mucin processing rather than mucin structure itself. There exists evidence for conservation of this pathway in mollusks as well, where a distinct N-acetylgalactosaminetransferase was characterized in snails and shown to differentially initiate glycolsation of various mucins, including MUC5AC, further suggesting that mucin processing may vary depending on the identity of the transferase and substrate (Taus et al. 2013). Together with the *mucin-5ac-like* gene identified in this study, these findings suggest that both mucus production and post-translational processing contribute to resistance against dermo infection.

Several additional loci implicate cellular signaling and apoptotic pathways in resistance to *P. marinus*. Significant markers were associated with genes encoding a TRAF-type zinc-finger protein and a mitogen-activated protein kinase kinase kinase (MAP3K). MAPK signaling has previously been implicated in molluscan hemocyte responses to pathogen exposure (Iakovleva et al. 2006). TRAF-family proteins function as key regulators of apoptosis and innate immune signaling (Lin et al. 2023) and are involved in transcriptional response to dermo (Chan et al. 2021). Although mollusks lack adaptive immunity, homologous signaling pathways regulate pathogen recognition, apoptosis, and inflammatory responses in invertebrates (Zhang et al. 2015). Variation in these signaling proteins may therefore influence the efficiency with which oysters detect infection and coordinate downstream immune responses.

Other implicated genes included several involved in cellular metabolism, including an ATP synthase that is responsible for ATP production in the mitochondria. This finding is consistent with differential expression of genes related to cellular metabolism in Eastern oysters infected by *P. marinus* (Chan et al. 2021, Proestou et al. 2023), suggesting regulation of cellular metabolism is important in the resistance of dermo, a wasting disease. Also, implicated are two genes involved in the maintenance of the extracellular matrix (ECM), which is often disrupted by *P. marinus*. Several other genes related to ECM function were also implicated in dermo resistance in another study (Wang et al. 2025). The involvement of genes related to ubiquitination in dermo response is also supported by transcriptomic data (Chan et al. 2021).

### Applications of Identified DRMs in Advancing Dermo Resistance

The DRMs identified in this study can accelerate selective breeding for dermo resistance through both marker-assisted selection and genomic selection. DRMs with relatively large and consistent effects, such as the *mucin-5AC-like* marker, could be incorporated into marker-assisted selection to prioritize favorable genotypes, while the larger set of small-effect DRMs may be more effectively incorporated into genomic selection approaches which capture their combined effects. Indeed, inclusion of these DRMs significantly improved the accuracy of nine different genomic prediction models trained using the dataset described in this study (Sun et al. 2026).

Currently, many aquaculture breeding programs cannot fully leverage genomic data due to the prohibitive cost of large-scale sequencing. However, smaller, targeted genotyping panels may significantly reduce costs while providing comparable accuracy and genetic gain to high-density arrays (Kriaridou et al. 2020, Fraslin et al. 2023, Wang et al. 2025, Delomas et al. 2026). Inclusion of DRMs into future SNP arrays or targeted genotyping panels could further increase representation of informative, low-frequency variants while enabling cost-effective selection in commercial breeding programs.

The identified DRMs also provide candidates for functional validation of biological mechanisms underlying dermo resistance. Because markers identified by GWAS are most likely linked to causal mutations, additional fine-mapping and quantitative trait loci studies need to be performed to specifically identify those causative variants. When causal variants are in strong LD with SNPs included on existing genotyping platforms, these markers may serve as cost-effective proxies for selection. Similar approaches are established in cattle breeding, where a SNP (rs109146371) genotyped using a commercially mass–produced chip is used to detect the presence of an unincluded SNP (rs109234250) that directly results in increased milk macronutrient concentration (Akiyama et al. 2025).

Validation of the DRMs using independent populations is important because both LD and marker effects may vary among genetic backgrounds and across environmental gradients. For example, in dairy cattle, some breeds exhibited near-perfect LD between the previously discussed SNPs, and others ranged widely (Strucken et al. 2010, Akiyama et al. 2025). Identifying causal variants and validating linked markers across independent breeding populations would ultimately provide more robust selection targets. Functional characterization of causal variants could also identify candidates for targeted genetic modification, providing additional opportunities to improve dermo resistance in Eastern oyster stocks.

## CONCLUSION

A multi-generational dataset and a two-stage GWAS approach with more inclusive QC criteria substantially improved the detection of markers associated with dermo resistance in *C. virginica*. The identification of 48 dermo-resistance markers, most characterized by deleterious rare alleles and small individual effects, supports the contention that dermo resistance is a polygenic trait with low-to-moderate heritability. The association between rare alleles and dermo susceptibility may be part of a general phenomenon where rare genetic variants play a critical role in shaping fitness. These results also demonstrate that conventional GWAS filtering may exclude low-frequency variants that likely contribute meaningfully to disease resistance and other complex traits in oysters. The identified dermo-resistance markers were associated with genes from diverse pathways, including innate immunity and metabolism. Most notably, the strongest association occurred within a *mucin-5ac-like* gene, suggesting mucus production is critical for dermo resistance. The identified DRMs have improved genomic prediction accuracy and provide strong candidates for advancing marker-assisted selection, genomic selection, targeted genotyping, and functional validation.

## DATA AVAILABILITY STATEMENT

Raw genotypic and phenotypic data generated by this study are available upon request to the corresponding author. All code used in this study is available at https://github.com/henrysun9074/gsAI/blob/main/analysis/GWASDarpaManuscript.R.

## ACKNOWLEDGMENTS

The authors thank the Center for Aquaculture Technology for their service of DNA extraction and SNP array genotyping.

## FUNDING STATEMENT

This study was supported by the Defense Advanced Research Projects Agency (DARPA) under Contract No. HR0011122C0136 and partly by the National Oceanic and Atmospheric Administration (NOAA), United States Department of Commerce through the Atlantic States Marine Fisheries Commission (award NA18NMF4720321). XG, DB, and JLP were also supported by USDA NIFA Hatch projects 1021665/NJ30401, 1009201/NJ32114, and LAB94509, respectively. The statements, opinions, findings, conclusions, and recommendations are those of the authors and do not necessarily reflect the views of NOAA, the Department of Commerce, DARPA or USDA. HS was supported by the NSF Graduate Research Fellowship under grant DGE-2139754.

## CONFLICT OF INTEREST STATEMENT

The authors declare no conflicts of interest.

## Notes

https://github.com/henrysun9074/gsAI/blob/main/analysis/GWASDarpaManuscript.R

### Competing Interest Statement

The authors have declared no competing interest.

### Summary of Updates

Supplementary Data File S1 was added; Figure 1 revised; references updated; typo in the author's name corrected.

## References 

Ahmad, S., Jose da Costa Gonzales, L., Bowler-Barnett, E. H., Rice, D. L., Kim, M., Wijerathne, S., Luciani, A., Kandasaamy, S., Luo, J., Watkins, X., Turner, E., Martin, M. J., & the UniProt Consortium. (2025). The UniProt website API: Facilitating programmatic access to protein knowledge. Nucleic Acids Research, 53(W1), W547–W553 10.1093/nar/gkaf394

Akiyama, Y., Ando, T., Nozaki, N., Arif, M., Ide, Y., Wang, S., & Miura, N. (2025). Strong linkage disequilibrium and proxy effect of PPP1R16A rs109146371 for DGAT1 K232A in Japanese Holstein cattle. Genes, 16(9), 1000. 10.3390/genes16091000

Allam, B., & Pales Espinosa, E. (2015). Mucosal immunity in mollusks. In Mucosal Health in Aquaculture (pp. 325–370). Academic Press. 10.1016/B978-0-12-417186-2.00012-1

Allam, B., & Raftos, D. (2015). Immune responses to infectious diseases in bivalves. Journal of Invertebrate Pathology, 131, 121–136. 10.1016/j.jip.2015.05.005

Allam, B., Carden, W. E., Ward, J. E., Ralph, G., Winnicki, S., & Espinosa, E. P. (2013). Early host-pathogen interactions in marine bivalves: Evidence that the alveolate parasite Perkinsus marinus infects through the oyster mantle during rejection of pseudofeces. Journal of Invertebrate Pathology, 113(1), 26–34. 10.1016/j.jip.2012.12.011

Altschul, S. F., Gish, W., Miller, W., Myers, E. W., & Lipman, D. J. (1990). Basic local alignment search tool. Journal of Molecular Biology, 215(3), 403–410. 10.1016/S0022-2836(05)80360-2

Anderson, C. A., Pettersson, F. H., Clarke, G. M., Cardon, L. R., Morris, A. P., & Zondervan, K. T. (2010). Data quality control in genetic case-control association studies. Nature Protocols, 5(9), 1564–1573. 10.1038/nprot.2010.116

Aquaculture | Economic Research Service. (n.d.). Retrieved April 16, 2026, from https://www.ers.usda.gov/topics/animal-products/aquaculture

ASReml Cookbook • ASReml Knowledge Base. (n.d.). ASReml Knowledge Base. Retrieved April 16, 2026, from https://asreml.kb.vsni.co.uk/knowledge-base/apiolaza-asreml-cookbook/

Audemard, C., Calvo, L. R., Paynter, K. T., Reece, K. S., & Burreson, E. M. (2006). Real-time PCR investigation of parasite ecology: In situ determination of oyster parasite Perkinsus marinus transmission dynamics in lower Chesapeake Bay. Parasitology, 132(6), 827–842. 10.1017/S0031182006009851

Bennett, E. P., Mandel, U., Clausen, H., Gerken, T. A., Fritz, T. A., & Tabak, L. A. (2012). Control of mucin-type O-glycosylation: A classification of the polypeptide GalNAc- transferase gene family. Glycobiology, 22(6), 736–756. 10.1093/glycob/cwr182

Bidegain, G., Powell, E. N., Klinck, J. M., Hofmann, E. E., Ben-Horin, T., Bushek, D., Ford, S. E., Munroe, D. M., & Guo, X. (2017). Modeling the transmission of Perkinsus marinus in the Eastern oyster Crassostrea virginica. Fisheries Research, 186, 82–93. 10.1016/j.fishres.2016.08.006

Boeckmann, B., Bairoch, A., Apweiler, R., Blatter, M.-C., Estreicher, A., Gasteiger, E., Martin, M. J., Michoud, K., O’Donovan, C., Phan, I., Pilbout, S., & Schneider, M. (2003). The SWISS-PROT protein knowledgebase and its supplement TrEMBL in 2003. Nucleic Acids Research, 31(1), 365–370. 10.1093/nar/gkg095

Bower, S. M. (2025). Synopsis of infectious diseases and parasites of commercially exploited shellfish: Perkinsus marinus (“Dermo” disease) of oysters. Fisheries and Oceans Canada. https://www.dfo-mpo.gc.ca/science/aah-saa/diseases-maladies/pmdoy-eng.html

Brookfield, J. F. Y. (1996). A simple new method for estimating null allele frequency from heterozygote deficiency. Molecular Ecology, 5(3), 453–455. 10.1046/j.1365-294X.1996.00098.x

Brown, B. L., Butt, A. J., Meritt, D., & Paynter, K. T. (2005). Evaluation of resistance to Dermo in eastern oyster strains tested in Chesapeake Bay. Aquaculture Research, 36, 1544– 1554. 10.1111/j.1365-2109.2005.01377.x

Butler, D. G., Cullis, B. R., Gilmour, A. R., Gogel, B. G., & Thompson, R. (2023). ASReml-R Reference Manual Version 4.2. VSN International Ltd., Hemel Hempstead, UK.

Burreson, E. M., & Ragone Calvo, L. M. (1996). Epizootiology of Perkinsus marinus disease of oysters in Chesapeake Bay, with emphasis on data since 1985. Journal of Shellfish Research, 15(1), 17–34.

Bushek, D., & Allen, S. K., Jr. (1996). Host-parasite interactions among broadly distributed populations of the eastern oyster Crassostrea virginica and the protozoan Perkinsus marinus. Marine Ecology Progress Series, 139, 127–141. 10.3354/meps139127

Bushek, D., Ford, S. E., & Chintala, M. M. (2002). Comparison of in vitro-cultured and wild- type Perkinsus marinus. III. Fecal elimination and its role in transmission. Diseases of Aquatic Organisms, 51(3), 217–225. 10.3354/dao051217

Bushek, D., Ford, S. E., & Burt, I. (2012). Long-term patterns of an estuarine pathogen along a salinity gradient. Journal of Marine Research, 70(2–3), 225–251. 10.1357/002224012802851968

Carnegie, R. B., Ford, S. E., Crockett, R. K., Kingsley-Smith, P. R., Bienlien, L. M., Safi, L. S., Whitefleet-Smith, L. A., & Burreson, E. M. (2021). A rapid phenotype change in the pathogen Perkinsus marinus was associated with a historically significant marine disease emergence in the eastern oyster. Scientific Reports, 11, 12872. 10.1038/s41598-021-92379-6

Carter, A. J. R., & Nguyen, A. Q. (2011). Antagonistic pleiotropy as a widespread mechanism for the maintenance of polymorphic disease alleles. BMC Medical Genetics, 12, 160. 10.1186/1471-2350-12-160

Casas, S., Walton, W., Chaplin, G., Rikard, S., Supan, J., & La Peyre, J. (2017). Performance of oysters selected for dermo resistance compared to wild oysters in northern Gulf of Mexico estuaries. Aquaculture Environment Interactions, 9, 169–180. 10.3354/aei00222

Chan, J., Wang, L., Li, L., Mu, K., Bushek, D., Xu, Y., Guo, X., Zhang, G., & Zhang, L. (2021). Transcriptomic response to Perkinsus marinus in two Crassostrea oysters reveals evolutionary dynamics of host-parasite interactions. Frontiers in Genetics, 12. 10.3389/fgene.2021.795706

Choi, K. S., Wilson, E. A., Lewis, D. H., Powell, E. N., & Ray, S. M. (1989). The energetic cost of Perkinsus marinus parasitism in oysters: Quantification of the thioglycollate method. Journal of Shellfish Research, 8, 125–131.

Cooperative Regional Oyster Selective Breeding (CROSBreed) Project. (2001). Potential of selected stocks for restoration and extensive planting. NOAA Oyster Disease Research Program.

Corbeil, R. R., & Searle, S. R. (1976). Restricted maximum likelihood (REML) estimation of variance components in the mixed model. Technometrics, 18(1), 31–38. 10.1080/00401706.1976.10489397

Cvijović, I., Good, B. H., & Desai, M. M. (2018). The effect of strong purifying selection on genetic diversity. Genetics, 209(4), 1235–1278. 10.1534/genetics.118.301058

Daniels, M. A., & Teixeiro, E. (2025). The NF-κB signaling network in the life of T cells. Frontiers in Immunology, 16. 10.3389/fimmu.2025.1559494

Dégremont, L., Garcia, C., & Allen, S. K. (2015). Genetic improvement for disease resistance in oysters: A review. Journal of Invertebrate Pathology, 131, 226–241. 10.1016/j.jip.2015.05.010

Dégremont, L., Lamy, J.-B., Pépin, J.-F., Travers, M.-A., & Renault, T. (2015). New insight for the genetic evaluation of resistance to Ostreid herpesvirus infection, a worldwide disease, in Crassostrea gigas. PLOS ONE, 10(6), e0127917. 10.1371/journal.pone.0127917

Dégremont, L., Nourry, M., & Maurouard, E. (2015). Mass selection for survival and resistance to OsHV-1 infection in Crassostrea gigas spat in field conditions: Response to selection after four generations. Aquaculture, 446, 111–121.

Delomas, T. A., Proestou, D. A., & Small, J. M. (2026). Evaluation of genomic selection to improve survival of eastern oysters infected with Perkinsus marinus. Frontiers in Genetics, 17. 10.3389/fgene.2026.1821653

Devlin, B., & Roeder, K. (1999). Genomic control for association studies. Biometrics, 55(4), 997–1004. 10.1111/j.0006-341X.1999.00997.x

Dickson, S. P., Wang, K., Krantz, I., Hakonarson, H., & Goldstein, D. B. (2010). Rare variants create synthetic genome-wide associations. PLOS Biology, 8(1), e1000294. 10.1371/journal.pbio.1000294

Duffy, M. A., & Forde, S. E. (2009). Ecological feedbacks and the evolution of resistance. Journal of Animal Ecology, 78(6), 1106–1112. 10.1111/j.1365-2656.2009.01568.x

Ernande, B., Clobert, J., McCombie, H., & Boudry, P. (2003). Genetic polymorphism and trade- offs in the early life-history strategy of the Pacific oyster, Crassostrea gigas (Thunberg, 1795): A quantitative genetic study. Journal of Evolutionary Biology, 16, 399–414. 10.1046/j.1420-9101.2003.00543.x

Ewart, J. W., & Ford, S. E. (1993). History and impact of MSX and Dermo diseases on oyster stocks in the northeast region (p. 8). Dartmouth, Massachusetts: Northeastern Regional Aquaculture Center, University of Massachusetts Dartmouth.

Fernández Robledo, J. A., Caler, E., Matsuzaki, M., Keeling, P. J., Shanmugam, D., Roos, D. S., & Vasta, G. R. (2011). The search for the missing link: A relic plastid in Perkinsus? International Journal for Parasitology, 41(12), 1217–1229. 10.1016/j.ijpara.2011.07.008

Fraslin, C., Robledo, D., Kause, A., & Houston, R. D. (2023). Potential of low-density genotype imputation for cost-efficient genomic selection for resistance to Flavobacterium columnare in rainbow trout (Oncorhynchus mykiss). Genetics Selection Evolution, 55, 59. 10.1186/s12711-023-00832-z

Gibson, G. (2012). Rare and common variants: Twenty arguments. Nature Reviews Genetics, 13(2), 135–145. 10.1038/nrg3118

Gignoux-Wolfsohn, S. A., Newcomb, M. S., Ruiz, G. M., & Lohan, K. M. P. (2021). Environmental factors drive the release of Perkinsus marinus from infected oysters. Parasitology, 148(5), 532–538. 10.1017/S0031182020002383

Gillis, J., & Pavlidis, P. (2013). Assessing identity, redundancy and confounds in Gene Ontology annotations over time. Bioinformatics, 29(4), 476–482. 10.1093/bioinformatics/bts727

Goddard, M. E., Hayes, B. J., & Meuwissen, T. H. E. (2010). Genomic selection in livestock populations. Genetics Research, 92(5–6), 413–421. 10.1017/S0016672310000613

Gómez-Chiarri, M., Warren, W. C., Guo, X., & Proestou, D. (2015). Developing tools for the study of molluscan immunity: The sequencing of the genome of the eastern oyster, Crassostrea virginica. Fish & Shellfish Immunology, 46(1), 2–4. 10.1016/j.fsi.2015.05.004

Guo, X. (2021). Genetics in shellfish culture. In S. E. Shumway (Ed.), Molluscan shellfish aquaculture: A practical guide (pp. 393–413). Essex, UK: 5M Books Ltd.

Guo, X. (2026a). Genetics and epigenetics in life history and reproduction: Bivalve molluscs. In M. K. Skinner (Ed.), Encyclopedia of reproduction (pp. 964–972). Elsevier.

Guo, X. (2026b). Overview of molluscan aquaculture and breeding. In S. Wang & Z. Bao (Eds.), Genetic breeding and aquaculture of shellfish (pp. 25–46). Elsevier.

Guo, X., Ford, S. E., DeBrosse, G., Smolowitz, R., & Sunila, I. (2003). Breeding and evaluation of eastern oyster strains selected for MSX, Dermo and JOD resistance. Journal of Shellfish Research, 22, 333–334.

Guo, X., Li, C., Wang, H., & Xu, Z. (2018). Diversity and evolution of living oysters. Journal of Shellfish Research, 37(4), 755–771. 10.2983/035.037.0407

Guo, X., Puritz, J. B., Wang, Z., et al. (2023). Development and evaluation of high-density SNP arrays for the Eastern oyster Crassostrea virginica. Marine Biotechnology, 25, 174–191. 10.1007/s10126-022-10191-3

Haldane, J. B. S. (1937). The effect of variation of fitness. The American Naturalist, 71(735), 337–349. 10.1086/280722

Harrang, E., Lapègue, S., Morga, B., & Bierne, N. (2013). A high load of non-neutral amino-acid polymorphisms explains high protein diversity despite moderate effective population size in a marine bivalve with sweepstakes reproduction. G3: Genes|Genomes|Genetics, 3(2), 333–341. 10.1534/g3.112.005181

He, Y., Yu, H., Bao, Z., Zhang, Q., & Guo, X. (2012). Mutation in promoter region of a serine protease inhibitor confers Perkinsus marinus resistance in the eastern oyster (Crassostrea virginica). Fish & Shellfish Immunology, 33(2), 411–417. 10.1016/j.fsi.2012.05.028

Hedgecock, D., & Pudovkin, A. (2011). Sweepstakes reproductive success in highly fecund marine fish and shellfish: A review and commentary. Bulletin of Marine Science, 87, 971–1002. 10.5343/bms.2010.1051

Iakovleva, N. V., Gorbushin, A. M., & Storey, K. B. (2006). Modulation of mitogen-activated protein kinases (MAPK) activity in response to different immune stimuli in haemocytes of the common periwinkle Littorina littorea. Fish & Shellfish Immunology, 21(3), 315– 324. 10.1016/j.fsi.2005.12.008

Impute. (n.d.). Bioconductor. Retrieved April 16, 2026, from http://bioconductor.org/packages/impute/

Itoïz, S., Metz, S., Derelle, E., Reñé, A., Garcés, E., Bass, D., Soudant, P., & Chambouvet, A. (2022). Emerging parasitic protists: The case of Perkinsea. Frontiers in Microbiology, 12. 10.3389/fmicb.2021.735815

Johansen, C. T., Wang, J., Lanktree, M. B., Cao, H., McIntyre, A. D., Ban, M. R., et al. (2010). Excess of rare variants in genes identified by genome-wide association study of hypertriglyceridemia. Nature Genetics, 42(8), 684–687. 10.1038/ng.628

Kachmar, M. L., Bergman, C., Schreier, H. J., Feild, G., Pagenkopp Lohan, K. M., Carnegie, R. B., Burge, C. A., & Gignoux-Wolfsohn, S. (2025). Spatio-temporal patterns of Perkinsus marinus infections are driven by a changing environment in the Chesapeake Bay. Diseases of Aquatic Organisms, 164, 111–127. 10.3354/dao03876

Koc, E. C., Burkhart, W., Blackburn, K., Moyer, M. B., Schlatzer, D. M., Moseley, A., & Spremulli, L. L. (2001). The large subunit of the mammalian mitochondrial ribosome. Journal of Biological Chemistry, 276(47), 43958–43969. 10.1074/jbc.M106510200

Kriaridou, C., Tsairidou, S., Houston, R. D., & Robledo, D. (2020). Genomic prediction using low density marker panels in aquaculture: Performance across species, traits, and genotyping platforms. Frontiers in Genetics, 11, 124. 10.3389/fgene.2020.00124

Kryukov, G. V., Pennacchio, L. A., & Sunyaev, S. R. (2007). Most rare missense alleles are deleterious in humans: Implications for complex disease and association studies. The American Journal of Human Genetics, 80(4), 727–739. 10.1086/513473

La Peyre, J. F., Casas, S. M., Richards, M., Xu, W., & Xue, Q. (2019). Testing plasma subtilisin inhibitory activity as a selective marker for dermo resistance in eastern oysters. Diseases of Aquatic Organisms, 133, 127–139. 10.3354/dao03344

La Peyre, J. F., Chu, F. L. E., & Vogelbein, W. K. (1995). In vitro interaction of Perkinsus marinus merozoites with eastern and Pacific oyster hemocytes. Developmental and Comparative Immunology, 19(4), 291–304. 10.1016/0145-305X(95)00017-N

Lamparter, D., Marbach, D., Rueedi, R., Kutalik, Z., & Bergmann, S. (2016). Fast and rigorous computation of gene and pathway scores from SNP-based summary statistics. PLoS Computational Biology, 12(1), e1004714. 10.1371/journal.pcbi.1004714

Lang, T., Hansson, G. C., & Samuelsson, T. (2007). Gel-forming mucins appeared early in metazoan evolution. Proceedings of the National Academy of Sciences, 104(41), 16209– 16214. 10.1073/pnas.0705984104

Lau, Y.-T., Gambino, L., Santos, B., Pales Espinosa, E., & Allam, B. (2018). Transepithelial migration of mucosal hemocytes in Crassostrea virginica and potential role in Perkinsus marinus pathogenesis. Journal of Invertebrate Pathology, 153, 122–129. 10.1016/j.jip.2018.03.004

Launey, S., & Hedgecock, D. (2001). High genetic load in the Pacific oyster Crassostrea gigas. Genetics, 159(1), 255–265. 10.1093/genetics/159.1.255

Lenz, T. L., Spirin, V., Jordan, D. M., & Sunyaev, S. R. (2016). Excess of deleterious mutations around HLA genes reveals evolutionary cost of balancing selection. Molecular Biology and Evolution, 33(10), 2555–2564. 10.1093/molbev/msw127

Li, J., & Ji, L. (2005). Adjusting multiple testing in multilocus analyses using the eigenvalues of a correlation matrix. Heredity, 95(3), 221–227. 10.1038/sj.hdy.6800717

Lima, A. K., Dhillon, H., & Dillman, A. R. (2022). ShK-domain-containing protein from a parasitic nematode modulates Drosophila melanogaster immunity. Pathogens, 11(10), 1094. 10.3390/pathogens11101094

Lin, M., Ji, X., Lv, Y., Cui, D., & Xie, J. (2023). The roles of TRAF3 in immune responses. Disease Markers, 2023, 7787803. 10.1155/2023/7787803

Liu, M., Fu, Q., Wang, P., Qiang, G., Luo, K., Chen, B., Luan, S., Kong, J., & Xu, P. (2026). Pedigree-assisted genotype imputation enables cost-effective genomic prediction in Penaeus vannamei. Scientific Reports. 10.1038/s41598-026-47716-y

Mackin, J. G. (1951). Histopathology of infection of *Crassostrea virginica* by *Dermocystidium marinum* Mackin, Owen and Collier. Bulletin of Marine Science of the Gulf and Caribbean, 1(1), 72–87.

Mackin, J. G., & Boswell, J. L. (1956). The life cycle and relationships of Dermocystidium marinum. Proceedings of the National Shellfisheries Association, 46, 112–115.

Marees, A. T., de Kluiver, H., Stringer, S., Vorspan, F., Curis, E., Marie-Claire, C., & Derks, E. M. (2018). A tutorial on conducting genome-wide association studies: Quality control and statistical analysis. International Journal of Methods in Psychiatric Research, 27(2), e1608. 10.1002/mpr.1608

Marion, S. B., & Noor, M. A. F. (2023). Interrogating the roles of mutation–selection balance, heterozygote advantage, and linked selection in maintaining recessive lethal variation in natural populations. Annual Review of Animal Biosciences, 11, 77–91. 10.1146/annurev-animal-050422-092520

Meuwissen, T. H., Hayes, B. J., & Goddard, M. E. (2001). Prediction of total genetic value using genome-wide dense marker maps. Genetics, 157(4), 1819–1829. 10.1093/genetics/157.4.1819

Morris, R. L., Akers, J. E., Casas, S., Geldard, J., Goad, A., Ghisalberti, M., Guo, X., Holland, R., Jin, H., Kibler, K. M., Kreeger, D., Lanham, B. S., La Peyre, J. F., Lowe, R. J., Lupton, C. E. M., Nassif, H., Mather, W., Pomeroy, A. W. M., Rothermel, E., Ruszala, M., Rupasinghe, M., Rikard, F. S., Riman, R. E., San Nicolas, R., Shinn, J. P., Shrestha, P., Sparks, E. L., Swearer, S. E., Taye, J., Temple, N. A., Thenuwara, M. N., Vien, P. B., Wang, Z., Williams, M. L., & Bushek, D. (2026). Reefense: Living shoreline mosaics can achieve ecological and engineering outcomes with interdisciplinary design. Proceedings of the National Academy of Sciences of the United States of America, 123(15), e2516197123. 10.1073/pnas.2516197123

NOAA Fisheries. (2024. Farmed eastern oysters. NOAA. Retrieved April 13, 2026, from https://www.fisheries.noaa.gov/species/eastern-oyster

Pales Espinosa, E., Corre, E., & Allam, B. (2014). Pallial mucus of the oyster Crassostrea virginica regulates the expression of putative virulence genes of its pathogen Perkinsus marinus. International Journal for Parasitology, 44(5), 305–317. 10.1016/j.ijpara.2014.01.006

Plough, L. V. (2016). Genetic load in marine animals: A review. Current Zoology, 62(6), 567– 579. 10.1093/cz/zow096

Plough, L. V., Shin, G., & Hedgecock, D. (2016). Genetic inviability is a major driver of type III survivorship in experimental families of a highly fecund marine bivalve. Molecular Ecology, 25(4), 895–910. 10.1111/mec.13524

Powell, E. N., Klinck, J. M., Guo, X., Hofmann, E. E., Ford, S. E., & Bushek, D. (2012). Can oysters Crassostrea virginica develop resistance to dermo disease in the field: The impediment posed by climate cycles. Journal of Marine Research, 70(2), 309–355. 10.1357/002224012802851896

Primmer, C. R., Papakostas, S., Leder, E. H., Davis, M. J., & Ragan, M. A. (2013). Annotated genes and nonannotated genomes: Cross-species use of Gene Ontology in ecology and evolution research. Molecular Ecology, 22(12), 3216–3241. 10.1111/mec.12309

Proestou, D. A., & Sullivan, M. E. (2020). Variation in global transcriptomic response to Perkinsus marinus infection among eastern oyster families highlights potential mechanisms of disease resistance. Fish & Shellfish Immunology, 96, 141–151. 10.1016/j.fsi.2019.12.001

Proestou, D. A., Sullivan, M. E., Lundgren, K. M., Ben-Horin, T., Witkop, E. M., & Hart, K. M. (2023). Understanding Crassostrea virginica tolerance of Perkinsus marinus through global gene expression analysis. Frontiers in Genetics, 14. 10.3389/fgene.2023.1054558

Puritz, J. B., Guo, X., Hare, M., He, Y., Hillier, L. W., Jin, S., Liu, M., Lotterhos, K. E., Minx, P., Modak, T., Proestou, D., Rice, E. S., Tomlinson, C., Warren, W. C., Witkop, E., Zhao, H., & Gomez-Chiarri, M. (2024). A second unveiling: Haplotig masking of the eastern oyster genome improves population-level inference. Molecular Ecology Resources, 24, e13801. 10.1111/1755-0998.13801

Ragone Calvo, L. M., Calvo, G. W., & Burreson, E. M. (2003). Dual disease resistance in a selectively bred eastern oyster, Crassostrea virginica, strain tested in Chesapeake Bay. Aquaculture, 220(1–4), 69–87. 10.1016/S0044-8486(02)00399-X

Ragone Calvo, L. M., Dungan, C. F., Roberson, B. S., & Burreson, E. M. (2003). Systematic evaluation of factors controlling Perkinsus marinus transmission dynamics in lower Chesapeake Bay. Diseases of Aquatic Organisms, 56(1), 75–86. 10.3354/dao056075

Rameez, R., Jahageerdar, S., Jayaraman, J., Chanu, T. I., Bangera, R., & Gilmour, A. (2022). Evaluation of alternative methods for estimating the precision of REML-based estimates of variance components and heritability. Heredity, 128(4), 197–208. 10.1038/s41437-022-00509-1

Rapp, K., Wei, S., Roberts, M., Yao, S., Fei, S. S., Gao, L., Ray, K., Wang, A., Godiah, R., & Han, L. (2024). Transcriptional profiling of mucus production in rhesus macaque endocervical cells under hormonal regulation. Biology of Reproduction, 111(5), 1045– 1055. 10.1093/biolre/ioae121

Ray, S. M., Mackin, J. G., & Boswell, J. L. (1953). Quantitative measurement of the effect on oysters of disease caused by *Dermocystidium marinum*. Bulletin of Marine Science of the Gulf and Caribbean, 3(1), 6–33.

Regional Shellfish Seed Biosecurity Program. (2026. About the program. Retrieved August 23, 2026, from https://rssbp.org/about-the-program/

Schwartz, L. C., Stauffer, B. A., Lavaud, R., La Peyre, M. K., Pandelides, A. F., McCarty, A. J., Small, J. M., & Plough, L. V. (2026). Genomic selection for low salinity tolerance in the eastern oyster Crassostrea virginica in Louisiana and Chesapeake Bay populations. Aquaculture, 619, 743812. 10.1016/j.aquaculture.2026.743812

Scott, L. R. (2025). Seasonal dynamics of an oyster pathogen during a period of climate change: Perkinsus marinus in Delaware Bay 1999–2024 [M.S. thesis, Rutgers, The State University of New Jersey, School of Graduate Studies]. ProQuest. https://www.proquest.com/docview/3257348049/abstract/92A7E07EFED409FPQ/1

Skol, A. D., Scott, L. J., Abecasis, G. R., & Boehnke, M. (2006). Joint analysis is more efficient than replication-based analysis for two-stage genome-wide association studies. Nature Genetics, 38(2), 209–213. 10.1038/ng1706

Smolowitz, R. (2013). A review of current state of knowledge concerning Perkinsus marinus effects on Crassostrea virginica (Gmelin) (the Eastern oyster). Veterinary Pathology, 50(3), 404–411. 10.1177/0300985813480806

Soniat, T. M. (1996). Epizootiology of Perkinsus marinus disease of eastern oysters in the Gulf of Mexico. Journal of Shellfish Research, 15(1), 35–43.

Soniat, T. M., Klinck, J. M., Powell, E. N., & Hofmann, E. E. (2006). Understanding the success and failure of oyster populations: Climatic cycles and Perkinsus marinus. Journal of Shellfish Research, 25(1), 83–93. 10.2983/0730-8000(2006)25%255B83%3AUTSAFO%255D2.0.CO;2

Soudant, P., Chu, E. F.-L., & Volety, A. (2013). Host–parasite interactions: Marine bivalve molluscs and protozoan parasites, Perkinsus species. Journal of Invertebrate Pathology, 114(2), 196–216. 10.1016/j.jip.2013.06.001

Strucken, E. M., Rahmatalla, S., De Koning, D.-J., & Brockmann, G. A. (2010). Haplotype analysis and linkage disequilibrium for DGAT1. Archives Animal Breeding, 53(3), 247– 255. 10.5194/aab-53-247-2010

Sullivan, M. E., & Proestou, D. A. (2021). Survival and transcriptomic responses to different Perkinsus marinus exposure methods in an eastern oyster family. Aquaculture, 542, 736831. 10.1016/j.aquaculture.2021.736831

Sun, H., Coyne, P., Wang, Z., Casas, S., La Peyre, J., Williams, M., Rikard, S., Bushek, D., Wong, J., & Guo, X. (2026). Multigenerational machine learning-based genomic prediction for dermo resistance in eastern oyster *Crassostrea virginica*. bioRxiv 2026.09.08.750100; doi: 10.64898/2026.09.08.750100

Supek, F., Bošnjak, M., Škunca, N., & Šmuc, T. (2011). REVIGO summarizes and visualizes long lists of Gene Ontology terms. PLOS ONE, 6(7), e21800. 10.1371/journal.pone.0021800

Sved, J. A., & Hill, W. G. (2018). One hundred years of linkage disequilibrium. Genetics, 209(3), 629–636. 10.1534/genetics.118.300642

Szklarczyk, D., Kirsch, R., Koutrouli, M., Nastou, K., Mehryary, F., Hachilif, R., Gable, A. L., Fang, T., Doncheva, N. T., Pyysalo, S., Bork, P., Jensen, L. J., & von Mering, C. (2023). The STRING database in 2023: Protein-protein association networks and functional enrichment analyses for any sequenced genome of interest. Nucleic Acids Research, 51(D1), D638–D646. 10.1093/nar/gkac1000

Tasumi, S., & Vasta, G. R. (2007). A galectin of unique domain organization from hemocytes of the eastern oyster (Crassostrea virginica) is a receptor for the protistan parasite Perkinsus marinus. The Journal of Immunology, 179(5), 3086–3098. 10.4049/jimmunol.179.5.3086

Taus, C., Lucini, C., Sato, T., Furukawa, K., Grabherr, R., & Staudacher, E. (2013). Expression and characterization of the first snail-derived UDP-N-acetyl-α-D- galactosamine:polypeptide N-acetylgalactosaminyltransferase. Glycoconjugate Journal, 30(9), 825–833. 10.1007/s10719-013-9486-6

The Gene Ontology Consortium. (2000). Gene Ontology: Tool for the unification of biology. Nature Genetics, 25, 25–29. 10.1038/75556

The UniProt Consortium. (2025). UniProt: The Universal Protein Knowledgebase in 2025. Nucleic Acids Research, 53(D1), D609–D617. 10.1093/nar/gkae1010

Thornton, D. J., & Sheehan, J. K. (2004). From mucins to mucus: Toward a more coherent understanding of this essential barrier. Proceedings of the American Thoracic Society, 1(1), 54–61. 10.1513/pats.2306016

Tian, E., Syed, Z. A., Edin, M. L., Zeldin, D. C., & Ten Hagen, K. G. (2023). Dynamic expression of mucins and the genes controlling mucin-type O-glycosylation within the mouse respiratory system. Glycobiology, 33(6), 476–489. 10.1093/glycob/cwad031

Tomczak, A., Mortensen, J. M., Winnenburg, R., Liu, C., Alessi, D. T., Swamy, V., Vallania, F., Lofgren, S., Haynes, W., Shah, N. H., Musen, M. A., & Khatri, P. (2018). Interpretation of biological experiments changes with evolution of the Gene Ontology and its annotations. Scientific Reports, 8(1), 5115. 10.1038/s41598-018-23395-2

Troyanskaya, O., Cantor, M., Sherlock, G., Brown, P., Hastie, T., Tibshirani, R., Botstein, D., & Altman, R. B. (2001). Missing value estimation methods for DNA microarrays. Bioinformatics, 17(6), 520–525. 10.1093/bioinformatics/17.6.520

Turner, S. D. (2014). qqman: An R package for visualizing GWAS results using Q-Q and Manhattan plots [Preprint]. bioRxiv. 10.1101/005165

U.S. Department of Agriculture, National Agricultural Statistics Service. (2019). 2018 Census of Aquaculture. https://www.nass.usda.gov/Publications/AgCensus/2017/Online_Resources/Aquaculture/

Uffelmann, E., Huang, Q. Q., Munung, N. S., de Vries, J., Okada, Y., Martin, A. R., Martin, H. C., Lappalainen, T., & Posthuma, D. (2021). Genome-wide association studies. Nature Reviews Methods Primers, 1, 59. 10.1038/s43586-021-00056-9

Vasta, G. R., Feng, C., Tasumi, S., Abernathy, K., Bianchet, M. A., Wilson, I. B. H., Paschinger, K., Wang, L.-X., Iqbal, M., Ghosh, A., Amin, M. N., Smith, B., Brown, S., & Vista, A. (2020). Biochemical characterization of oyster and clam galectins: Selective recognition of carbohydrate ligands on host hemocytes and Perkinsus parasites. Frontiers in Chemistry, 8, 98. 10.3389/fchem.2020.00098

Venditti, D. A., Steele, C. A., Ayers, B. S., & McCormick, J. L. (2022). How long can dead fish tell tales? Effects of time, tissue, preservation, and handling on genotyping success. Northwest Science, 95(3–4), 337–349. 10.3955/046.095.0309

Visscher, P. M., Wray, N. R., Zhang, Q., Sklar, P., McCarthy, M. I., Brown, M. A., & Yang, J. (2017). 10 years of GWAS discovery: Biology, function, and translation. The American Journal of Human Genetics, 101(1), 5–22. 10.1016/j.ajhg.2017.06.005

Wang, J., Gao, J., Sheng, X., Tang, X., Xing, J., Chi, H., & Zhan, W. (2024). Teleost Muc2 and Muc5ac: Key guardians of mucosal immunity in flounder (Paralichthys olivaceus). International Journal of Biological Macromolecules, 277, 134127. 10.1016/j.ijbiomac.2024.134127

Wang, S., Peatman, E., Liu, H., Bushek, D., Ford, S. E., Kucuktas, H., Quilang, J., Li, P., Wallace, R., & Wang, Y. (2010). Microarray analysis of gene expression in eastern oyster (Crassostrea virginica) reveals a novel combination of antimicrobial and oxidative stress host responses after dermo (Perkinsus marinus) challenge. Fish & Shellfish Immunology, 29(6), 921–929. 10.1016/j.fsi.2010.07.035

Wang, Z., Casas, S., Peyre, J. L., Rikard, S., Williams, M. L., Tarnecki, A., Bushek, D., & Guo, X. (2025). Genomic selection for dermo resistance in the eastern oyster Crassostrea virginica: Production and laboratory testing of F1 generation. Journal of Shellfish Research, 44(1), 75–87. 10.2983/035.044.0108

Wiggans, G. R., & Carrillo, J. A. (2022). Genomic selection in United States dairy cattle. Frontiers in Genetics, 13, 994466. 10.3389/fgene.2022.994466

Yin, X., & Hedgecock, D. (2021). Overt and concealed genetic loads revealed by QTL mapping of genotype-dependent viability in the Pacific oyster Crassostrea gigas. Genetics, 219(4), iyab165. 10.1093/genetics/iyab165

Yu, Z., & Guo, X. (2006). Identification and mapping of disease-resistance QTLs in the eastern oyster, Crassostrea virginica Gmelin. Aquaculture, 254(1–4), 160–170. 10.1016/j.aquaculture.2005.10.016

Zhang, L., Li, L., Guo, X., Litman, G. W., Dishaw, L. J., & Zhang, G. (2015). Massive expansion and functional divergence of innate immune genes in a protostome. Scientific Reports, 5, 8693. 10.1038/srep08693

